# Tex2 is required for lysosomal functions at TMEM55-dependent ER membrane contact sites

**DOI:** 10.1101/2022.07.12.499833

**Authors:** Yuanjiao Du, Weiping Chang, Lin Deng, Wei-Ke Ji

## Abstract

ER tubules form and maintain membrane contact sites (MCSs) with late endosomes/lysosomes (LE/lys). The molecular composition and cellular functions of these MCSs are poorly understood. Here we find that Tex2, an SMP domain-containing lipid transfer protein conserved in metazoen and yeast, is a tubular ER protein, and enriches at ER-LE/lys MCSs dependent on TMEM55, phosphatases that convert PI(4,5)P2 to PI5P on LE/lys. We show that the Tex2-TMEM55 interaction occurs between a N-terminal region of Tex2 and a catalytic motif in PTase domain of TMEM55. The Tex2-TMEM55 interaction can be regulated by endosome-resident type 2 PI4K activities. Functionally, Tex2 knockout results in severe defects in lysosomal digestive capacity and blocked autophagic flow, as well as an aberrant accumulation of PI3P on the LE/lys membranes. These defects can be substantially rescued by wild type Tex2 other than a lipid transfer-defective Tex2 mutant, indicating an important role of lipid transfer in these processes. Together our data identify Tex2 as a tubular ER protein that resides at TMEM55-depedent ER-LE/lys MCSs required for lysosomal functions.

## Introduction

The endoplasmic reticulum (ER) is organized into a continuous intracellular membrane network, consisting of the nuclear envelope, flattened sheets, and interconnected tubules that spread over the cytosol(Baumann and Walz, 2001; Bian et al., 2011; English and Voeltz, 2013; Levine and Rabouille, 2005). ER tubules are considered as main sites for the synthesis of majority of lipids (Borgese et al., 2006). Newly synthesized lipids at the ER are then transfered to other organelles, through both vesicular and nonvesicular transport pathways. Although vesicular lipid transport mediates the bulk transport of many lipids, increasing lines of evidence suggest that lipid-transfer proteins (LTPs) mediated lipid transport at membrane contact sites (MCSs) is the major transport route for certain lipid types (Holthuis and Levine, 2005; Joshi et al., 2017; Levine, 2004; Levine, 2005; Prinz et al., 2020; Wong et al., 2017; Wong et al., 2019). MCSs are defined as cytosolic gaps of 10 – 30 nm between one organelle and the other organelles (Lebiedzinska et al., 2009; Levine, 2004).

The majority of early endosomes and all late endosomes and lysosomes (hereafter referred collectively to as LE/lys) maintain MCSs with ER tubules (Friedman et al., 2013; Zajac et al., 2013), which play critical roles in diverse cellular processes, including modulating endosome maturation and positioning, lipid composition, and fission during cargo sorting (Friedman et al., 2013; Gao et al., 2022; Hoyer et al., 2018; Jongsma et al., 2016; Raiborg et al., 2015; Rocha et al., 2009; Rowland et al., 2014; Wu and Voeltz, 2021). However, the molecular composition and cellular functions of these MCSs are poorly understood.

Tex2, is a synaptotagmin-like mitochondrial-lipid-binding (SMP) domain-containing LTP that resides on the ER. Tex2 is conserved in metazoen and yeast. Nvj2p, the Tex2 homolog in yeast, relocalizes from contacts between the ER and other organelles to and increases ER–Golgi contacts upon ER stress. Nvj2p may directly transfer ceramide from the ER to the Golgi complex destined for spingolipid synthesis, and prevents the buildup of toxic amounts of ceramides(Liu et al., 2017).

Tex2 is recently suggested to be present at ER-LE/lys MCSs along with another SMP-containing protein PDZD8 in worms (Jeyasimman et al., 2021). However, the mechanisms underlying the localization of Tex2 to these MCSs remain elusive. More importantly, cellular functions of Tex2-mediated MCSs remain largely unknown in mammals. In this study, we find that Tex2 is a tubular ER protein, and can be recruited to ER-LE/lys MCSs by TMEM55. The TMEM55-mediated recruitment of Tex2 is regulated by the activities of type 2 PI4Ks, which are resident on endosomal membranes and contribute to the generation of PI4P on endosomal membranes. Tex2 is required for endosome maturation and lysosomal digestive capacities in a lipid transfer-activity dependent manner. Together our data identify Tex2 as a tubular ER protein that may transport lipids at TMEM55-depedent ER-LE/lys MCSs essential for lysosomal functions.

## Results

### Tex2 is a tubular ER protein

E-Syts, a group of SMP domain-containing proteins, transfer a range of glycerophospholipids between the ER and the plasma membrane (PM) (Schauder et al., 2014). To identify unknown proteins that may interact with E-Syt1, we performed GFP-Trap assays in HEK293 cells transiently expressing GFP-E-Syt1 followed by mass spectrometry (MS). After removal of those proteins co-immunoprecipitated (coIPed) by GFP alone, we found a SMP domain-containing protein named testis expressing protein 2 (Tex2) that strongly interested us owing to its potential lipid transfer activity.

To begin with, we sought to explore the localization of Tex2 and its relations to E-Syt1. We confirmed the Tex2–E-syt1 interaction in GFP-Trap assays using GFP-Tex2 as a bait in HEK293 cells. The GFP-trap assays showed the Halo tagged E-Syt1 was coIPed with GFP-Tex2 (Fig. S1A). Since E-Syt1 is an integral ER protein, and is enriched at ER–PM MSCs upon ER calcium depletion (Bian et al., 2018; Giordano et al., 2013), we asked whether Tex2 colocalized with E-Syt1 at these MCSs by live-cell confocal microscopy. GFP-Tex2 partially colocalized with Halo-E-Syt1 on the ER under normal condition (Fig. S1B; top; S1C). Halo-E-Syt1 was enriched on the discrete ER microdomains, likely representing the ER-PM MCSs, upon thapsigargin (TG) treatment. However, GFP-Tex2 was not specifically enriched at these sites (Fig. S1B; bottom; S1C), suggesting that Tex2 may not function at ER-PM MCSs.

One of the most noticeable features was that GFP-Tex2 exclusively localizes to ER tubules other than sheets (Fig. 1A), whereas Halo-E-Syt1 was evenly distributed on both tubules and sheets of the ER (Fig. S1B). To avoid potential artifacts of overexpression, we labeled endogenous Tex2 with monomeric GFP (A206K) at its N terminus using Crispr-Cas9 in HeLa cells (GFP-Tex2-KI), in which ∼50% of endogenous Tex2 was labeled by GFP (Fig. S2A, B). Consistently, endogenous Tex2 is exclusively localized to ER tubules (Fig. 1B).

**Fig. 1.**
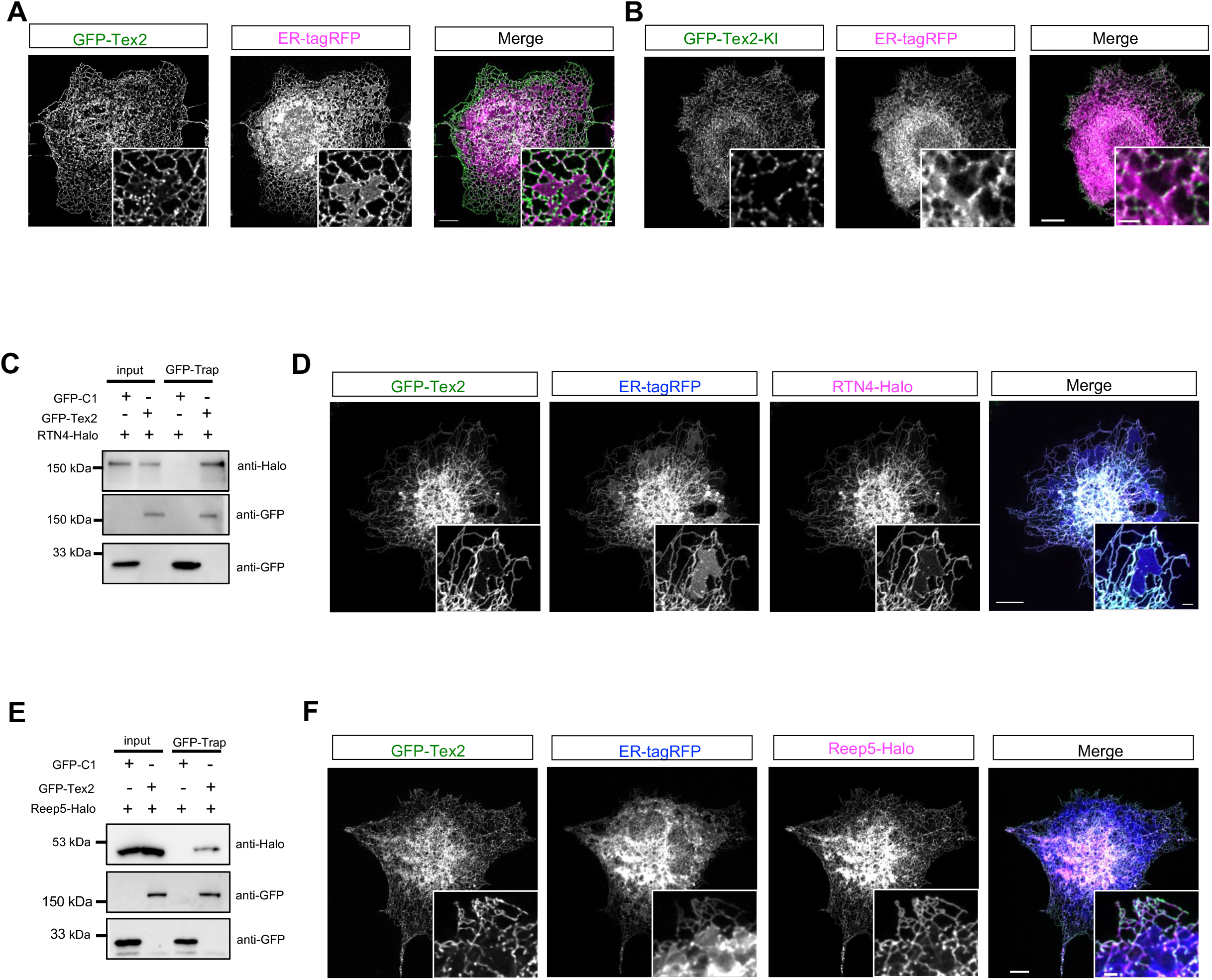
Tex2 is a tubular ER protein. **A.** Representative images of a live COS7 cell expressing exogenous GFP-Tex2 (green) and ER-tagRFP (magenta, an ER luminal marker) with insets. **B.** Representative images of a live GFP-Tex2-KI HeLa cell expressing endogenous GFP-Tex2 (green) and ER-tagRFP (magenta) with insets. **C.** GFP-Trap assays demonstrate an interaction between GFP-Tex2 and RTN4-Halo in COS7 cells. **D.** Representative images of a live COS7 cell expressing GFP-Tex2 (green), RTN4-Halo (magenta), and ER-tagRFP (blue) with insets. **E.** GFP-Trap assays from COS7 cells demonstrate an interaction between GFP-Tex2 and Reep5-Halo **F.** Representative images of a live COS7 cell expressing GFP-Tex2 (green), Reep5-Halo (magenta), and ER-tagRFP (blue) with insets. Scale bar, 10μm in the whole cell images and 2μm in the insets in (A, B, D, & F).

The formation of ER tubule is tightly controlled by a group of proteins, including Reep5 (Chen et al., 2021), Reticulon 4 (RTN4) (Voeltz et al., 2006), Atlastin-1 (ATL1) (Wang et al., 2016), and ARL6IP1 (Yamamoto et al., 2014). We next explore the relations between Tex2 and these ER-tubule shaping proteins. The GFP-trap assays showed that Tex2 interacts with RTN4 (Fig. 1C), and Reep5 (Fig. 1E), but interact with ATL-1 (Fig. S3A) and ARL6IP1 (Fig. S3C) to a much less extent. Meanwhile, live-cell confocal microscopy showed that GFP-Tex2 colocalized with RTN4 (Fig. 1D) and REEP5 (Fig. 1F) on the ER tubules, but only partially colocalized with other ER-shaping proteins (Figs. S3B, D; S1C). We then asked whether Tex2 is recruited to ER tubules by these ER shaping proteins. Suppression of these ER-shaping proteins by small interfering RNAs did not abolish tubular ER localization of Tex2, as revealed by live-cell confocal microscopy (Fig.S3E-J), suggesting that Tex2 may target tubular ER of its own.

Therefore, we dissected the Tex2 protein (Fig. 2A), and found that a N-terminal region containing a TM domain (Tex2-NT; residues 1–517) of Tex2 was sufficient for its ER tubule localization (Fig. 1B), but a smaller region only containing the TM domain (Tex2-TM; residues 475–517) failed to exclusively target ER tubules (Fig. 1C). Consistently, Tex2 without the NT region (Tex2-Δ1-473) reduced its specific localization (Fig. 1D, G), suggesting that the NT region is sufficient and required for Tex2 targeting to ER tubules. Notably, a portion of Tex2-Δ1-473 appeared to re-localize to some discrete ER regions, likely the ER-PM MCSs (Fig. 1D). We further found that the residues 277-517 of NT is sufficient to target ER tubules (Fig. 1E, G), and Tex2 with a deletion of this region (Tex2-Δ277-473) substantially reduced its specific targeting to tubular ER (Fig. 1F, G).

**Fig. 2.**
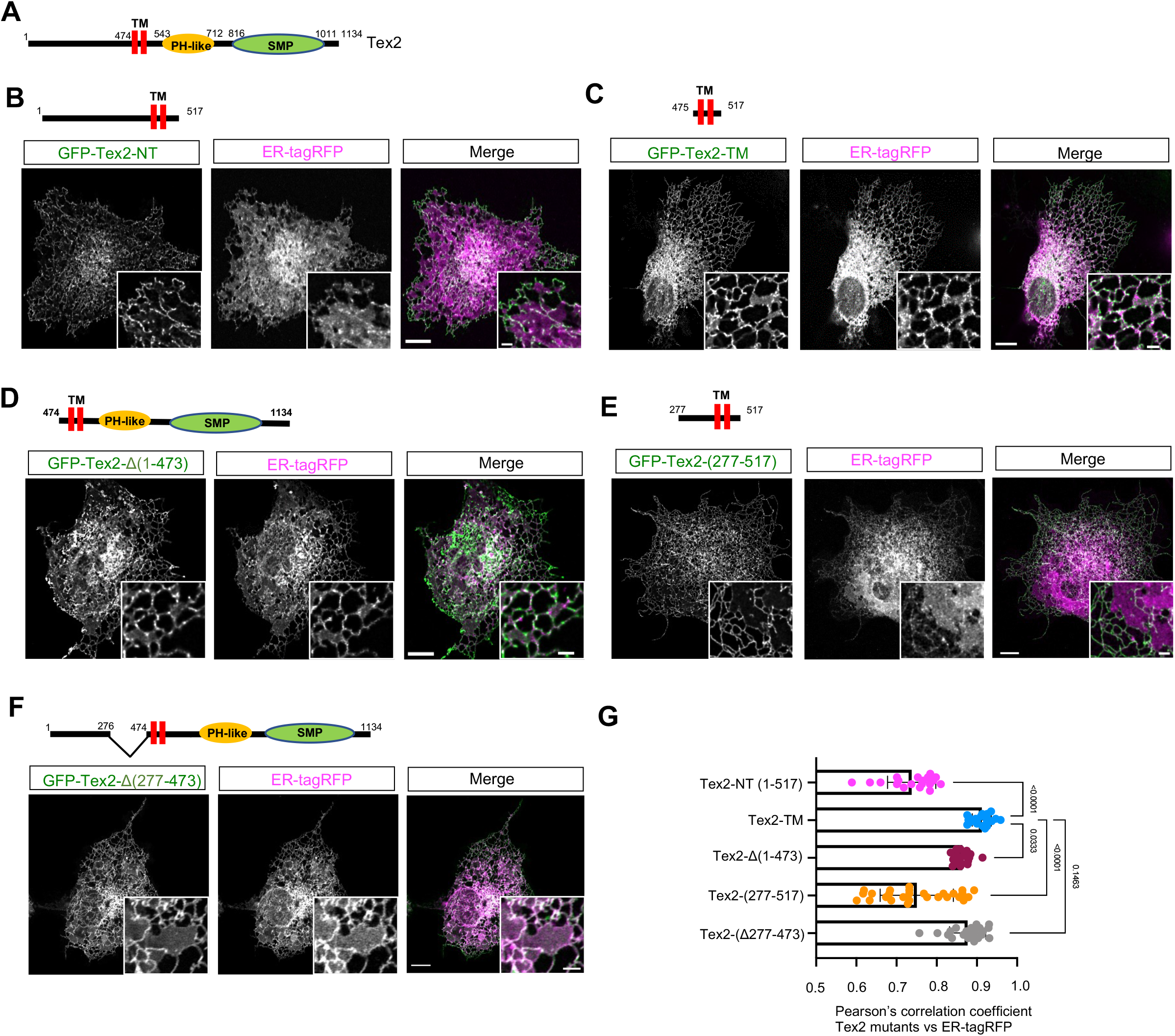
A N-terminal region adjacent to TM is required for Tex2 targeting tubular ER. **A**. Domain organization of Tex2. **B-F**. Representative images of COS7 cells expressing either GFP-Tex2-TM (green; residues 474-517; **B**), GFP-Tex2-1-517aa (green; **C**), GFP-Tex2-Δ1-473 (green; **D**), GFP-Tex2-(277-517) (green; **E**) or GFP-Tex2-Δ277-473 (green; **F**), along with ER-tagRFP (magenta) with insets. **G.** Pearson’s correlation coefficient of Tex2 proteins vs ER-tagRFP; GFP-Tex2-TM (22 cells); GFP-Tex2-1-517aa (19 cells); GFP-Tex2-Δ1-473 (23 cells); GFP-Tex2-(277-517) (26 cells); and GFP-Tex2-Δ277-473 (21 cells) in more than 3 independent experiments. Ordinary one-way ANOVA with Tukey’s multiple comparisons test. Mean ± SD. Scale bar, 10μm in the whole cell images and 2μm in the insets in (B-F, H).

Next, we investigated whether Tex2 is able to promote the formation of ER tubules. Overexpression of Climp63-Halo promoted the formation of ER sheet at cell periphery (Fig. S4A), consistent with a reported role of Climp63 in the formation of ER sheets (Shibata et al., 2010), which appeared to be countered upon co-expression of GFP-Tex2 (Fig. S4B), to a similar extent compared to the effects of RTN4 overexpression (Fig. S4C), a well-studied ER tubule-forming protein (Wang et al., 2016). Interestingly, Crispr-Cas9-mediated Tex2 KO appeared not to substantially affect the tubular ER network at periphery (Fig. S4D; S2C, D), suggesting a redundant role of these tubular ER-resident proteins in the formation and/or maintenance of the tubular ER network. Collectively, our results suggested that Tex2 exclusively targets tubular ER via the region (residues 277-474) adjacent to the TM.

### TMEM55B recruits Tex2 to ER-LE/Lys MCSs

As a potential LTP resident on ER tubules, Tex2 likely functions at ER-associated MCSs. To identify the type of ER-associated MCSs that Tex2 may function, we sought to identify its adaptor on the other organelle by coIP-MS using GFP-Tex2 as bait. To this end, we identified TMEM55B, a phosphatase that converts PI(4,5)P_2_ to PI5P on LE/lys as a novel Tex2-interacting protein (Fig. 3A). Consistently, we found that E-Syt1 and the tubular ER proteins, including RTN4, REEP5, RTN3, RTN1, and ARL6IP1 were identified in our MS analysis (Fig. 3A). Notably, TMEM55B did not rank at the top of our list. We reason that Tex2 was mainly on ER tubules without great enrichments at ER MCSs under normal condition in which our coIP-MS was performed, and suggested that Tex2-TMEM55B interactions may be subjected to regulation. Supporting our MS results, GFP-trap assays confirmed the interactions between GFP-Tex2 and Halo-TMEM55B (Fig. 3B). Importantly, live-cell confocal microscopy demonstrated that overexpression of Halo-TMEM55B greatly recruited GFP-Tex2 to another organelle that clustered at perinuclear regions (Fig. 3C, box1), whereas GFP-Tex2 was mainly localized to the entire ER tubules in cells without Halo-TMEM55B, with a minor enrichment at perinuclear regions (Fig. 3C, box2). As a control, Halo-TMEM55B failed to recruit GFP-E-Syt1 to perinuclear regions (Fig. 3D), suggesting a specific recruitment of Tex2 by TMEM55B.

**Fig. 3.**
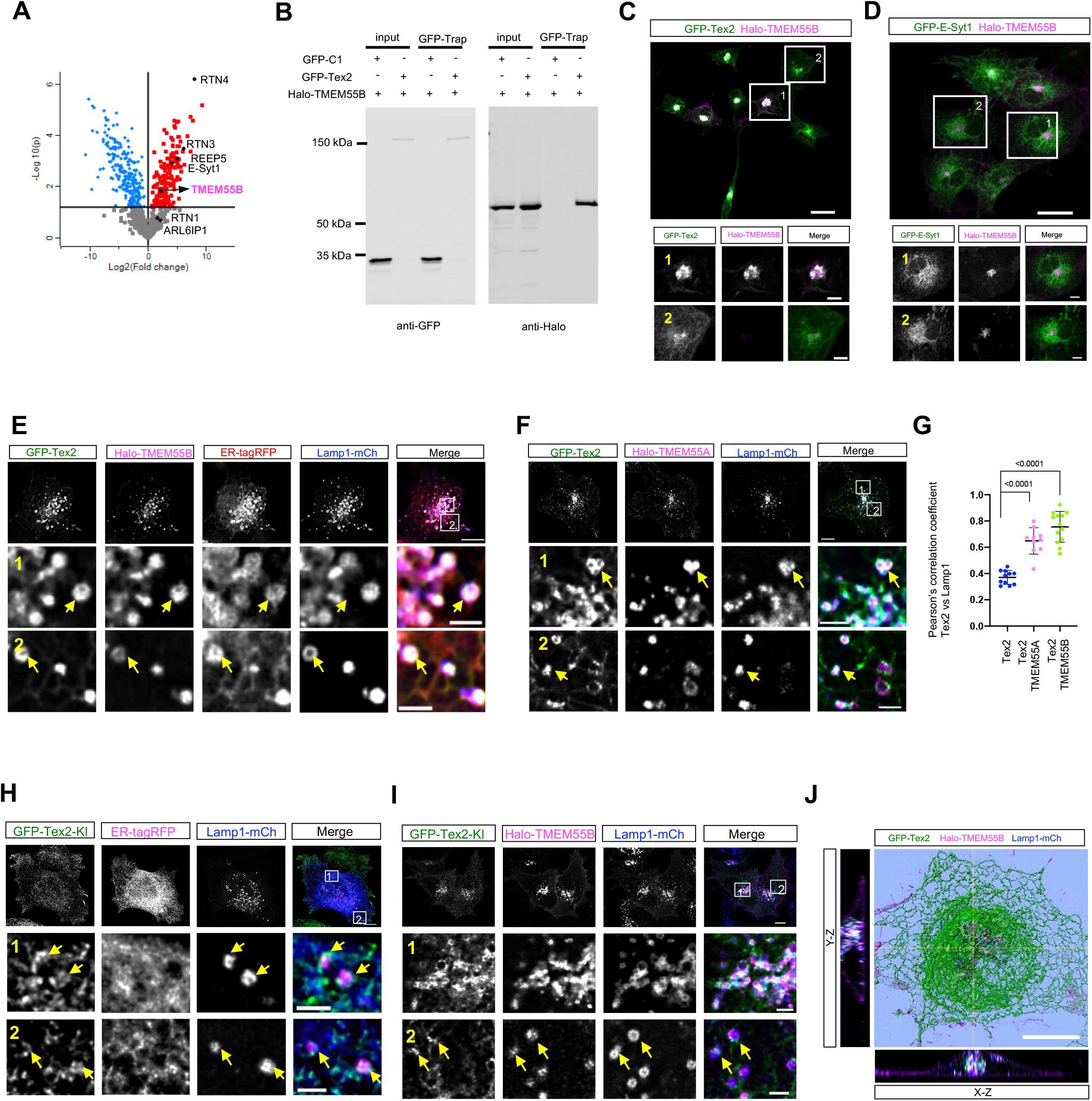
Identification of TMEM55B as a Tex2 adaptor on LE/Lys. **A.** Volcano plot of protein candidates coIPed with Tex2 in HEK293 cells. **B.** GFP-Trap assays demonstrate an interaction between GFP-Tex2 and Halo-TMEM55B in COS7 cells. **C-D**. Representative wide-field images of live COS7 cells expressing either GFP-Tex2 (green; C) or GFP-E-syt1 (green; D) and Halo-TMEM55B with two boxed regions showing at the bottom. **E-F**. Representative images of a live COS7 cell expressing GFP-Tex2 (green), Halo-TMEM55B (magenta; **E**) or Halo-TMEM55A (magenta; **F**), ER-tagRFP (red), and Lamp1-mCh (blue; a LE/lys marker) with two boxed regions showing at the bottom. **G**. Pearson’s correlation coefficient of Tex2 vs Lamp1 in absence of TMEM55 (11 cells) or upon TMEM55A (10 cells) or TMEM55B (13 cells) overexpression (> 3 independent experiments). Ordinary one-way ANOVA with Tukey’s multiple comparisons test. Mean ± SD. **H-I**. Representative live-cell images of a GFP-Tex2-KI (green) cell without (**H**) or with expressing Halo-TMEM55B (**I**) and Lamp1-mCh (blue) with two boxed regions at the bottom. Yellow arrows denote the specific enrichment of GFP-Tex2-KI at LE/lys extensively contacting the ER. **J**. Representative 3D rendering of a COS7 cell expressing GFP-Tex2 (green), Halo-TMEM55B (magenta), and Lamp1-mCh (blue) with y-z projection to the left and x-z projection to the right. Scale bar, 10μm in the whole cell images and 2μm in the insets in (C, D, E, F, H & I).

Next, we examined the type of the other organelle in Tex2-resident ER MCSs by live-cell confocal microscopy. We found that GFP-Tex2, Halo-TMEM55B and a LE/lys marker (Lamp1-mCh) were co-localized, with the ER (marked by a luminal ER marker ER-tagRFP) being strongly enriched at these sites (Fig. 3E), indicating that TMEM55B recruits Tex2 to ER-LE/lys MCSs.

TMEM55A and TMEM55B are phosphoinositide 4-phosphatases that dephosphorylate the D4 position of PI(4,5)P2 mainly on LE/lys membranes (Ungewickell et al., 2005). Human TMEM55A and TMEM55B share 51% identity in amino acid sequences. Both isozymes contain a CX_5_R motif in their phosphatase domains and two putative TM domains at the C-terminal. Next, we asked whether TMEM55A could recruit Tex2 by live-cell confocal microscopy as well. Halo-TMEM55A strongly recruited GFP-Tex2 to TMEM55A-positive LE/lys membranes (Fig. 3F), to a similar extent as TMEM55B (Fig. 3G). Since only TMEM55B was identified in our MS analysis, we focused on TMEM55B thereafter in this study.

Next, we confirmed the recruitment of endogenous Tex2 by TMEM55B in GFP-Tex2-KI cells. Live-cell confocal images showed that, though endogenous GFP-Tex2 was mainly distributed over tubular ER network, a small but significant portion of endogenous GFP-Tex2 was enriched at regions adjacent to LE/lys (Fig. 3H), likely ER-LE/lys MCSs, in absence of exogenous TMEM55B. Remarkably, upon Halo-TMEM55B expression, endogenous Tex2 was substantially enriched at TMEM55B-labeled LE/lys membranes (Fig. 3I), suggesting that TMEM55B greatly promoted the recruitment of endogenous GFP-Tex2 to the contacts.

We then used 3-dimension (3D) rendering of z-stacks through high-resolution live-cell microscopy to examine the localizations of GFP-Tex2 and Halo-TMEM55B relative to LE/lys. Tex2 was specifically enriched on TMEM55B-positive LE/lys at perinuclear (Fig. 3I) in 3D, as revealed by co-localization analysis based on x-z and y-z projections of 3D rendering.

Of note, we observed that LE/lys were substantially confined to perinuclear regions upon TMEM55B overexpression (Fig. S5A, B), in accord with a reported role of TMEM55B in promoting the retrograde trafficking of LE/lys (Willett et al., 2017). The phenotype could be allievated, to some extent, by co-expression of Tex2 and TMEM55B, whereas co-expression of Tex2 without the NT (Tex2-Δ1-517) with TMEM55B had no effect. In addition, another Tex2 mutant with a deletion of a PH-like domain (PH) (Tex2-ΔPH) was still able to allievate the phenotype resulted from TMEM55B overexpression to a similar extent as the wild type (WT) Tex2 (Fig. S5B), suggesting a role of Tex2-NT for interacting with TMEM55B. In addition, time-lapse video analysis showed that TMEM55B/Tex2-postive LE/lys preferentially underwent retrograde transport (Fig. S5D). Interestingly, we observed that the Tex2-labled ER membranes were stably assocaited with TMEM55B-positive LE/lys during the retrograde transport (Fig. S5D), which was rarely observed in cells (Fig. S5C). Collectively, these results indicate that TMEM55B acts as adaptor on LE/lys membranes to recruit ER-resident Tex2 to ER-LE/lys MCSs, and these MCSs may be involved in the regulation of the retrograde trafficking of LE/lys.

Given a specific role of Nvj2p, the yeast homolog of Tex2, at ER-Golgi MCSs upon ER stress, we explored whether Tex2 could be recruited to ER-Golgi MCSs upon TG-induced ER stress by live-cell confocal microscopy. The induction of ER stress was confirmed by a increase in BiP level (Fig. S6A), a marker of ER stress. Upon ER stress, endogenous GFP-Tex2 was still localized to the entire ER tubule network, and was likely not redirected to the Golgi apparatus upon either 2 h (Fig. S6B; middle) or 12 h treatment of TG (Fig. S6C; top), similar to untreated conditions (Fig. S6B; top; Fig. S6D). In addition, endogenous GFP-Tex2 was still substantially recruited to Halo-TMEM55B-positive LE/lys upon ER stress (Fig. S6B; bottom; Fig. S6C; bottom; Fig. S6E). Together, these results suggest that Tex2 may not be directly involved in TG-induced ER stress. However, it should be noted Tex2 may play important roles in other types of cellular stress, which is not identified in this study.

### The N-terminal region of Tex2 is responsible for interacting with TMEM55B

Next, we dissected the Tex2 protein to investigate how Tex2 interacted with TMEM55B. Live-cell microscopy showed that Tex2-ΔTM was mainly cytosolic in absence of TMEM55B (Fig. 4A; left), but a significant portion of this mutant was recruited to Halo-TMEM55B-positive LE/lys (Fig. 4A; right; 4F). Another Tex2 mutant with a deletion of both the TM and PH domain (Tex2-ΔTM-ΔPH) was still able to be recruited to TMEM55B-postitive LE/lys, similar to WT Tex2 (Fig. S7A, H). Consistently, a Tex2 mutant without the PH domain (Tex2-ΔPH) was recruited to TMEM55B-positive LE/lys, as revealed by the wrapping of Tex2-labeled ER tubules around LE/lys (Fig. S7B, H). These results suggested that the PH is not required for the Tex2-TMEM55B interactions. A PH domain is often involved in protein targeting to PIs-enriched membranes, including LE/lys membranes (Lemmon, 2007). Notably, PIPs strip assays using purified PH domain of Tex2 showed that Tex2-PH preferentially bound PI3P and PI4P, but bound, to a less extent, to PI5P, PI3,4P_2_, PI3,5P_2_, or PI4,5P_2_ (Fig. S7C). Since the formation of Tex2-mediated MCSs have been examined under condtions where TMEM55B is overexpressed in this study, it is possible that the PH domain may facilitate the recruitment of Tex2, independent of TMEM55B, to other membranes, for example, the PM by binding to PI(4,5)P2, under other conditions.

**Fig. 4.**
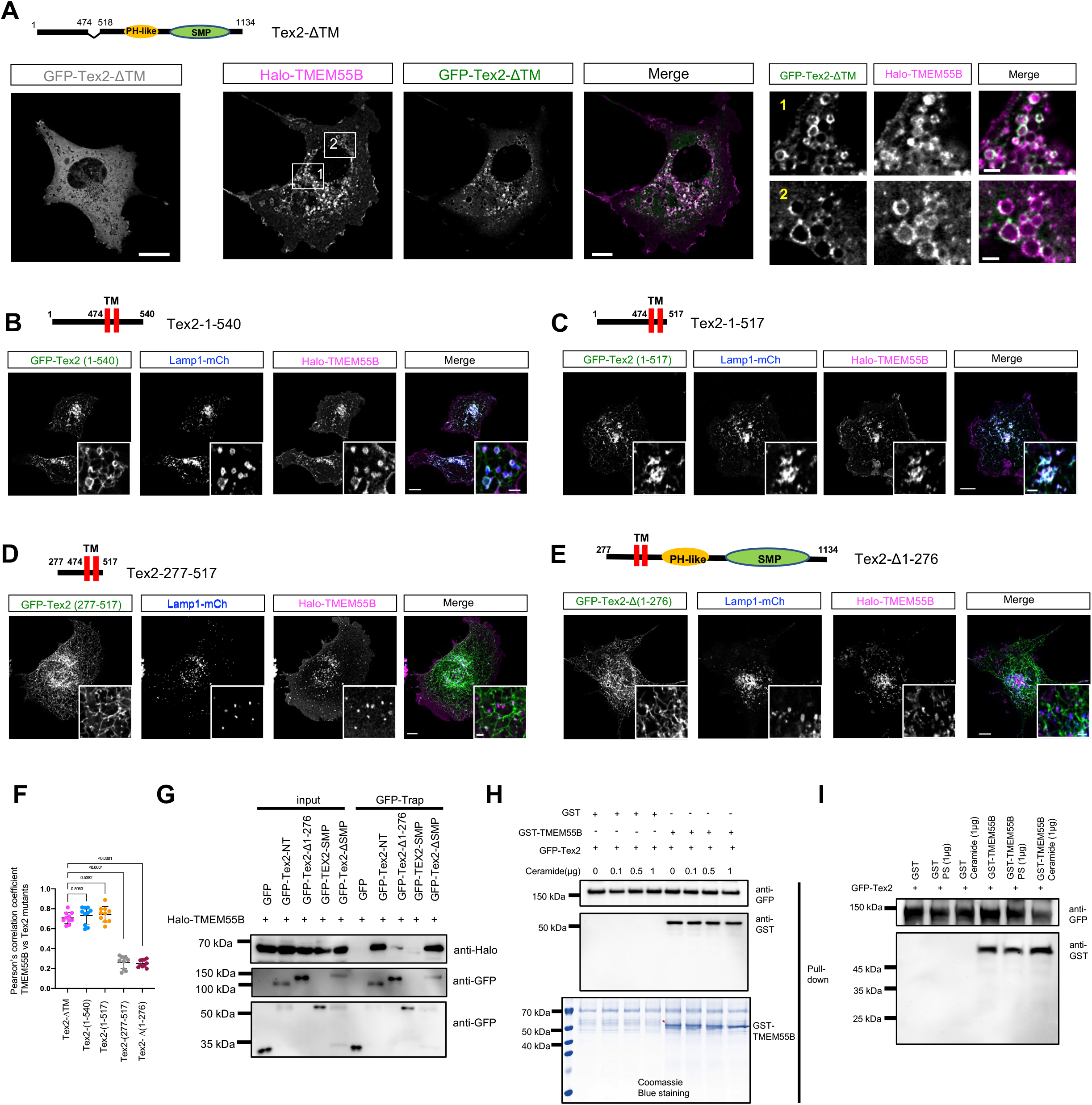
The N-terminal region of Tex2 is responsible for interacting with TMEM55B. **A**. Left: representative images of COS7 cells expressing GFP-Tex2-ΔTM (green) alone; middle: representative whole-cell images of a COS7 cell expressing GFP-Tex2-ΔTM (green), Halo-TMEM55B (magenta), and Lamp1-mCh (blue); right: two enlarged images from the two boxed regions in the middle panel. **B-E**. Representative images of COS7 cells expressing either GFP-Tex2-NT-1 (green; **B**), GFP-Tex2-NT-2 (green; **C**), GFP-Tex2 (277-517) (green; **C**) or GFP-Tex2-Δ (1-276) (green; **D**), along with Halo-TMEM55B (magenta), Lamp1-mCh (blue) with insets. **F**. Pearson’s correlation coefficient of TMEM55B vs Tex2 mutants; Tex2-ΔTM (10 cells); Tex2-1-540 (10 cells); Tex2-1-517 (10 cells); Tex2-277-517 (10 cells); and Tex2-Δ1-276 (10 cells) in more than 3 independent experiments. Ordinary one-way ANOVA with Tukey’s multiple comparisons test. Mean ± SD. **G**. GFP-Trap assays demonstrate interactions between Halo-TMEM55B and Tex2 mutants in COS7 cells. **H**. Pulldown assays using GFP-Tex2 bound on GFP-Trap beads and purified GST-TMEM55B demonstrate a direct interaction in a ceramide-independent manner. **I**. As in **(H)**, pulldown assays using GFP-Tex2 bound on GFP-Trap beads and purified GST-TMEM55B demonstrate a direct interaction in a PS-independent manner. Scale bar, 10μm in the whole cell images and 2μm in the insets in (A-E).

Next, we find that Tex2 without the SMP domain (Tex2-ΔSMP) could be recruited to TMEM55B-positive LE/lys, but to a slightly less extent compared to the WT-Tex2 (Fig. S7D, H). Consistently, the SMP domain alone (Tex2-SMP) was cytosolic even upon TMEM55B overexpression (Fig. S7E, H). These results suggested that the SMP domain may not be required for the Tex2-TMEM55B interactions.

Importantly, live-cell confocal miscroscopy showed that the residues 1-540 of Tex2 can be substantially recruited by TMEM55B (Fig. 4B, F). Further dissection on the NT demonstrated that the Tex2-NT (Tex2-1-517) (Fig. 4C, F), but not Tex2-277-517 (Fig. 4D, F), was sufficient for the recruiment. Consistently, Tex2 with a deletion of the residues 1-276 (Tex2-Δ1-276) failed to be recruited by TMEM55B (Fig. 4E, F), suggesting an essential role of the residues 1-276 for Tex2-TMEM55B interactions. Notably, we found that neither the residues 1-276 nor residues 1-474 of Tex2-NT could be recruited by TMEM55B (Fig. S7F, G, H), suggesting that these two regions were required but not sufficient for the recruitment. Together, these data indicated that the Tex2-NT (1-517) is the minimal functional module for the recruitment by TMEM55B.

Next, we sought to confirm the Tex2-TMEM55B interactions by GFP-Trap assays. In accord with the live-cell microscopy results, GFP-Tex2-NT was strongly coIPed with Halo-TMEM55B, whereas the level of Halo-TMEM55B coIPed by GFP-Tex2-Δ1-276 was greatly reduced (Fig. 4G), confirming a critical role of Tex2-NT in the Tex2-TMEM55B interactions. Consistently, Tex2-SMP could not be coIPed with Halo-TMEM55B (Fig. 4G), whereas Tex2-ΔSMP coIPed with Halo-TMEM55B in a similar level as Tex2-NT (Fig. 4G), confirming that Tex2-NT is responsible for interacting with TMEM55B.

We further asked whether Tex2 directly interact with TMEM55B by in vitro pulldown assays. At this time we were unable to produce purified full-length Tex2 or Tex2-NT in sufficient quantities for in vitro pull-down assays. Alternatively, we used GFP-Trap assays to pellet endogenous GFP-Tex2 from HeLa GFP-Tex2-KI using high-salt (500 mM NaCl) lysis buffer. After rigorous washing to remove proteins that could co-pellet with GFP-Tex2 under high-salt conditions, GFP-Tex2 beads were incubated with purified Glutathione S-transferase (GST) tag alone, or GST-TMEM55B, respectively. Indeed, western blots showed that GFP-Tex2, bound to GST-TMEM55B but not GST tag (Fig. 4H). This result suggests that GFP-Tex2 directly binds GST-TMEM55B. In addition, the interaction between GFP-Tex2 and GST-TMEM55B appeared not to be affected by an additon of Ceramide or PS (Figs. 4H and 4I), two lipid species that were shown to be bound by Tex2 later in this study (Fig. 7).

### A catalytic motif of TMEM55B is required for recruiting Tex2

Next, we sought to understand the molecular mechanisms underlying the recruitment of Tex2 by TMEM55B through dissections of the TMEM55B protein (Fig. 5A). Remarkably, live-cell microscopy showed that a TMEM55B truncation without C-terminal TM domains (TMEM55B-ΔTM) was cytosolic (Fig. 5B), but was greatly recruited to ER tubules upon the expression of GFP-Tex2 (Fig. 5C, H), indicating a reverse recruitment of cytosolic TMEM55B-ΔTM to the ER by GFP-Tex2. Another TMEM55B mutant without a region containing residues 1-74 (TMEM55B-ΔNT) still interacted with Tex2 to a similar extent as WT TMEM55B, as revealed by co-localization between GFP-Tex2 and this mutant on LE/lys (Fig. 5D, H). However, a truncation only containing the TM domain of TMEM55B (TMEM55B Δ1-163) failed to recruit GFP-Tex2 (Fig. 5E, H). Importantly, another mutant with a deletion of phosphatase domain (TMEM55B-ΔPTase) could target LEs, but failed to recruit GFP-Tex2 (Fig. 5F, H). These lines of evidence suggest that the PTase domain is critical for the recruitment of Tex2 by TMEM55B. We next investigated whether the recruitment of Tex2 is dependent on the highly conserved catalytic motif CX_5_R of the PTase domain. A mutant with a deletion of the CX_5_R motif (TMEM55B-ΔCX_5_R) compelely lost its ability to recruit Tex2 to LE/lys (Fig. 5G, H), indicating that the CX_5_R motif is required for interacting with Tex2. Collectively, our data indicated that the Tex2-TMEM55B interaction is dependent on the CX_5_R motif in the PTase domain of TMEM55B, and suggested that the reaction catalyzed by TMEM55B may be coupled with its interaction with Tex2.

**Fig. 5.**
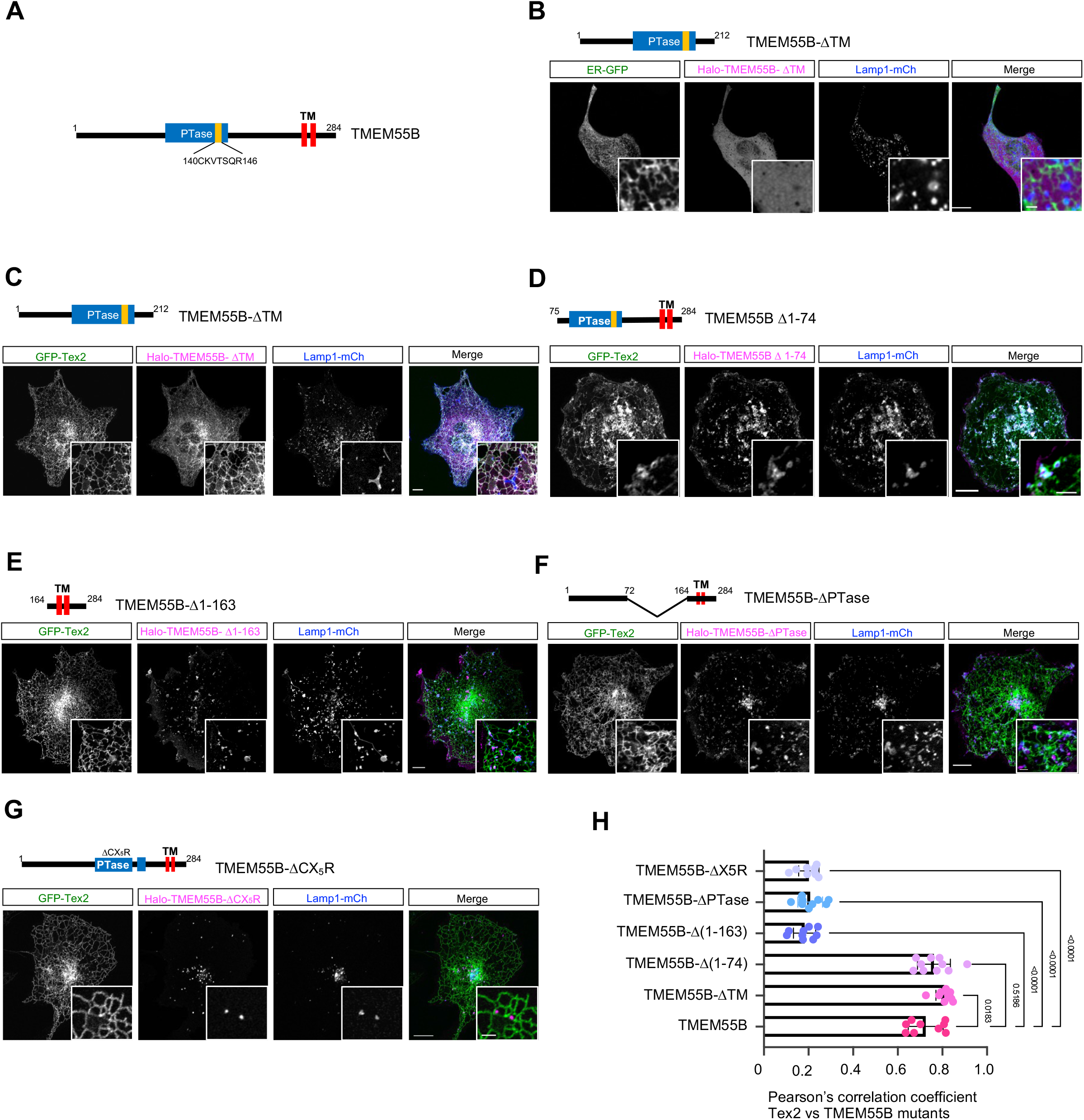
The catalytic motif CX_5_R of TMEM55B is responsible for interacting with Tex2. **A**. Domain organization of TMEM55B. **B**. Representative images of a COS7 cell expressing Halo-TMEM55B-ΔTM (red), ER-GFP (green), and Lamp1-mCh (blue) with insets. **C-G**. Representative images of COS7 cells expressing either Halo-TMEM55B-ΔTM (red; **C**), Halo-TMEM55B-ΔNT (red;75-284aa; **D**), Halo-TMEM55B-TM (red; **E**), Halo-TMEM55B-ΔPTase (red; **F**), and Halo-TMEM55B-ΔX5R (red; **G**). **H**. Pearson’s correlation coefficient of TMEM55B proteins vs Tex2; Halo-TMEM55B-ΔTM in absence of Tex2 (9 cells); and Halo-TMEM55B-ΔTM (9 cells); Halo-TMEM55B-Δ1-74 (10 cells); Halo-TMEM55B-Δ1-163 (9 cells); Halo-TMEM55B-ΔPTase (9 cells); Halo-TMEM55B-ΔCX_5_R (9 cells) upon Tex2 overexpression in 3 independent experiments. Ordinary one-way ANOVA with Tukey’s multiple comparisons test. Mean ± SD. Scale bar, 10μm in the whole cell images and 2μm in the insets in (**B-G**).

### The regulation of the Tex2-TMEM55B interaction by PI4KII activities

Next, we asked whether and how the Tex2-TMEM55B interaction was regulated. Importantly, we found that co-expression of PI4KIIα or PI4KII*Δ*, PI kinases that convert PI to PI4P on the membranes of LE/lys (Balla et al., 2002), significantly hampered the recruitment of GFP-Tex2 to LE/lys membranes by TMEM55B (Fig. 6A, B,& F; yellow arrows indicate diminished Tex2 enrichments at PI4KII-positive/TMEM55B-positive LE/lys; red arrows denoting Tex2 enrichments at PI4KII-negative/TMEM55B-positive LE/lys), whereas co-expression of the kinase-dead mutant PI4KIIα-W359A (Zhou et al., 2014) did not impair the TMEM55B-mediated recuitment of Tex2 (Fig. 6C, F), indicating that the activities of PI4KIIα play an important role in the regulation of Tex2-TMEM55 interactions. In addition, the co-localization between Tex2 and PI4KIIα-W359A were significantly higher than that of Tex2 and WT PI4KIIα (Fig. 68G), further supporting that the activities of PI4KII inhibits the recruitment of Tex2 to ER-LE/lys MCSs. Moreover, we found that the co-expression of PI4KIIα or PI4KII*Δ* did not substantially impair the recruitment of Tex2-NT by TMEM55B (Fig. 6D, F), suggesting a potential regulatory module at the C-terminal region of Tex2. Indeed, the recruitment of GFP-Tex2-ΔPH by TMEM55B was not significantly affacted by the co-expression of PI4KIIα or PI4KII*Δ* (Fig. 6E, F), suggesting that the PH domain may be involved in the regulation step. Consistently, the co-localization between Tex2-NT or Tex2-ΔPH and PI4KIIs were not substantially altered upon the loss of kinase activities of PI4KIIα (Fig. 6G). Given that purified Tex2-PH bound PI4P, PI(3,4)P_2_, and PI(4,5)P_2_ in vitro (Fig. S7C), it is tempting to speculate that the transient binding of these PIPs to the PH domain of Tex2 regulate the interaction with TMEM55B. Since PI4KIIs are responsible for the generation of PI4P pool on the LE/lys membrane, which can be further converted to other PIPs, such as PI(3,4)P_2_ and PI(4,5)P_2_, and thus it is still unclear which PIPs are responsible for the regulation of Tex2-TMEM55 interactions.

**Fig. 6.**
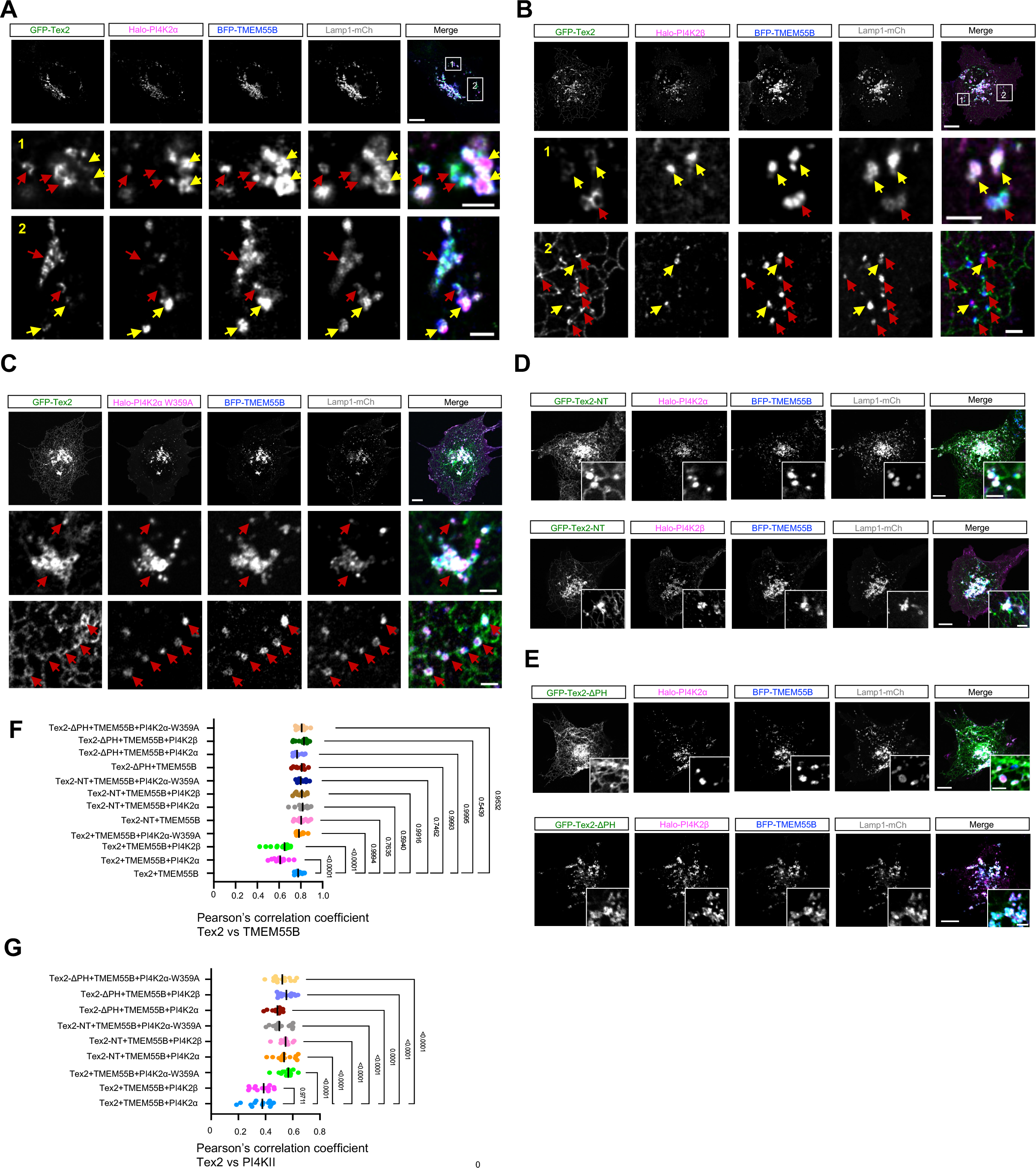
The regulation of the Tex2-TMEM55B interaction by PI4KII activities. **A-C**. Representative images of COS7 cells expressing either Halo-PI4KIIα (magenta; **A**), Halo-PI4KIIβ (magenta; **B**) or kinase-dead PI4KIIα mutant Halo-PI4KIIα-W359A (magenta; **C**), along with GFP-Tex2 (green), Lamp1-mCh (grey) and BFP-TMEM55B (blue) with two boxed regions showing at bottom. Yellow arrows indicate reduced Tex2 enrichments at PI4KII-positive/TMEM55B-positive LE/lys, while red arrows denote Tex2 enrichments at PI4KII-negative/TMEM55B-positive LE/lys. **D-E**. Representative images of COS7 cells expressing either GFP-Tex2-NT (green; **D**) or GFP-Tex2-ΔPH (green; **E**) along with Halo-PI4KIIs (magenta), Lamp1-mCh (grey) and BFP-TMEM55B (blue) with insets. **F**. Pearson’s correlation coefficient of Tex2 vs TMEM55B in cells expressing either empty vector (10 cells), Halo-PI4KIIα (13 cells), Halo-PI4KIIβ (14 cells), or Halo-PI4KIIα-W359A (13 cells). Pearson’s correlation coefficient of Tex2-NT vs TMEM55B in cells expressing either empty vector (14 cells), Halo-PI4KIIα (12 cells), Halo-PI4KIIβ (11 cells) or Halo-PI4KIIα-W359A (13 cells). Pearson’s correlation coefficient of Tex2-ΔPH vs TMEM55B in cells expressing either empty vector (12 cells), Halo-PI4KIIα (10 cells), Halo-PI4KIIβ (13 cells), or Halo-PI4KIIα-W359A (12 cells) in 3 independent assays. Ordinary one-way ANOVA with Tukey’s multiple comparisons test. Mean ± SD. **G**. Pearson’s correlation coefficient of Tex2 vs PI4KIIs in cells expressing either Halo-PI4KIIα (13 cells), Halo-PI4KIIβ (14 cells), or Halo-PI4KIIα-W359A (13 cells). Pearson’s correlation coefficient of Tex2-NT vs TMEM55B in cells expressing either Halo-PI4KIIα (12 cells), Halo-PI4KIIβ (11 cells) or Halo-PI4KIIα-W359A (13 cells). Pearson’s correlation coefficient of Tex2-ΔPH vs TMEM55B in cells expressing either Halo-PI4KIIα (10 cells), Halo-PI4KIIβ (13 cells), or Halo-PI4KIIα-W359A (12 cells) in 3 independent assays. Ordinary one-way ANOVA with Tukey’s multiple comparisons test. Mean ± SD. Scale bar, 10μm in the whole cell images and 2μm in the insets in (A-E).

### Tex2 binds glycerophospholipids and ceramides

Given that the SMP is a lipid-transfer domain, we asked whether the SMP domain of Tex2 might solubilize lipids, a prerequisite for a lipid transport function, by in vitro lipids-binding assays. Purified Tex2-SMP co-migrated with nitrobenzoxadiazole (NBD)-labeled glycerophospholipids [phosphatidylcholine (PC), phosphatidylserine (PS), phosphatidylethanolamine (PE)] and sphingolipids [ceramide (Cer)], but not cholesterol, as assessed by native gel electrophoreses (Fig. 7A, B). Among these lipids, the SMP domain preferentially bound to PC, PS and ceramide (Fig. 7C).

**Fig. 7.**
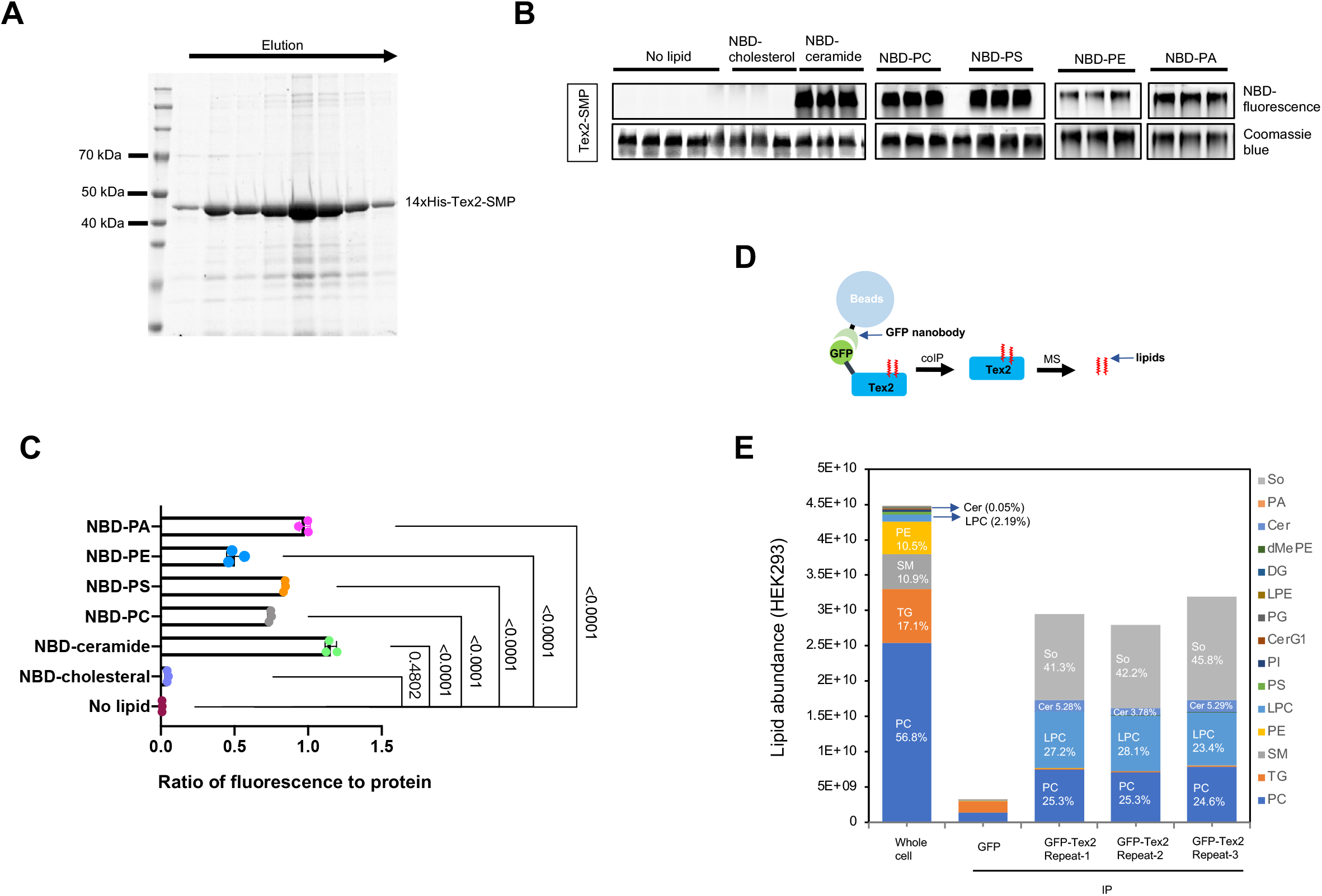
Tex2 binds glycerophospholipids and ceramides. **A.** Coomassie blue staining of purified 14xHis-Tex2-SMP. **B.** In vitro lipid-binding assays for Tex2-SMP. Purified Tex2-SMP was incubated with NBD-tagged lipids and examined by native PAGE. Phospholipids, visualized by their fluorescence, comigrated with protein, visualized by Coomassie blue staining. **C.** Ratio of fluorescence of Tex2-SMP-bound lipids to protein level. Ordinary one-way ANOVA followed by Tukey’s multiple comparisons test. Mean ± SD. **D.** Schematic cartoon of non-targeting lipidomic analysis of endogenous GFP-Tex2. **E.** Quantification of lipids bound to endogenous GFP-Tex2, from three independent assays. The lipid composition of total membranes in HEK293 cells were shown (Gao et al., 2022).

Next, we sought to identify lipid species bound by full-length GFP-Tex2 in HEK293 cells. Lipid species associating with GFP-Tex2 were assessed by non-targeted lipidomics using liquid chromatography-tandem mass spectrometry (LC-MS/MS) with rigorous washes of the protein before lipid analysis, according to the protocol used in our previous study (Fig. 7D) (Gao et al., 2022). Remarkably, GFP-Tex2 mainly associated with LPC/PC (∼50%) and sphingolipids [sphingosine (So) and Cer] (∼50%) (Fig. 7E). Taking into consideration the lipid composition of total membranes in HEK293 cells, PC accounted for ∼57% of the total phsopholipids, whereas spingolipids only accounts for less than 15% (Fig. 7E) (Gao et al., 2022), it is plausible that GFP-Tex2 may preferentially bind sphingolipids in cells. It should be noted that Tex2 may bind phosphatidylinositol phosphates (PIPs) in cells, but PIPs was too low to be detectable in our assays due to the low abundance of PIs in cells.

### Tex2 is required for lysosomal functions in a lipid transfer-dependent manner

We sought to explore the cellular functions of Tex2. Given that Tex2 acts at ER-LE/lys MCSs, we began to assess its roles on these two organelles. We showed that the localization of Tex2 was not affected by the TG-induced ER stress (Fig. S6B, C). Consistently, the initiation of ER stress response appeared not to be influenced by Tex2 KO, as revealed by the BiP level before and after TG stimulation (Fig. S6A). On the other hand, we explored the impacts of Tex2 KO on the functions of LE/lys by a lysosomal function sensor mApple-Lamp1-phLuorin (Fig. 8A). In control cells, very few LE/lys were marked by phLuorin fluoresence (Fig. 8B; left; E), indicating that majority of LE/lys were functional. Strikingly, two independent Tex2-KO clones, however, resulted in a remarkable increase in the percentage of phLuorin-positive LE/lys (Fig. 8B; right; E), which is similar to cells treated with Bafilomycin A1 (BafA1) (Fig. 8B; left), a potent v-ATPase inhibitor that blocks lysosomal functions.

**Fig.8.**
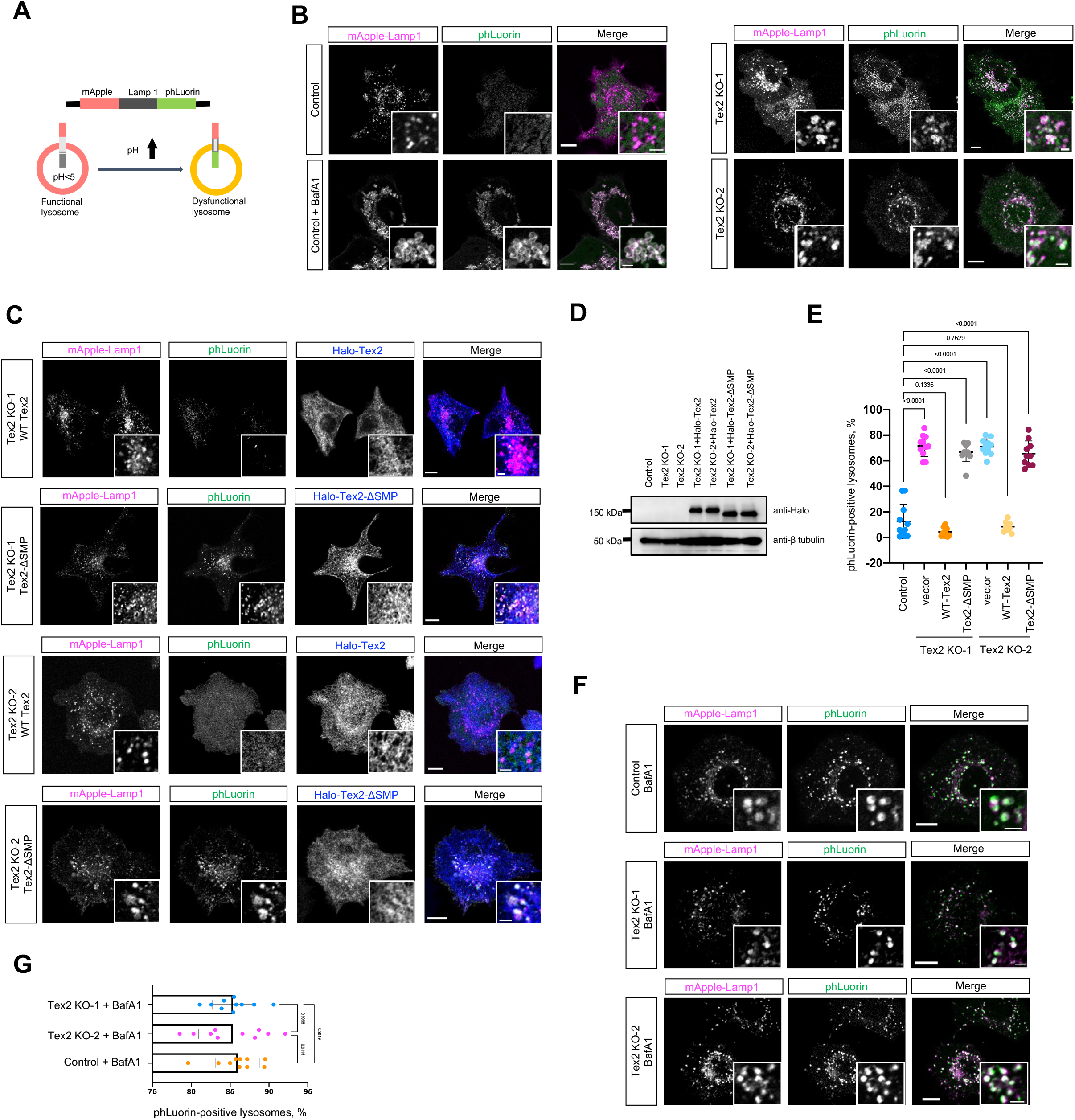
Tex2 is required for lysosomal functions in a lipid transfer-dependent manner. **A.** Schematic cartoon of mApple-Lamp1-phLuorin. **B.** Representative images of control (top), control cells treated with BafA1 (400nM; 2h; bottom), or two HeLa Tex2-KO clones (right), expressing mApple-Lamp1-phLuorin (phLuorin in green; mApple-Lamp1 in magenta) with insets. **C.** Representative images of two HeLa Tex2-KO clones expressing either Halo-Tex2 (blue) or Halo-Tex2-ΔSMP (blue) and mApple-Lamp1-phLuorin with insets. **D.** Western blots demonstrate the level of Halo-Tex2 and Halo-Tex2-ΔSMP in rescue experiments. **E.** Percentage of phLuorin-positive lysosomes in control (11 cells), Tex2 KO-1 (11 cells), Tex2 KO-1 resuced by Halo-Tex2 (10 cells), Tex2 KO-1 resuced by Halo-Tex2-ΔSMP (10 cells), Tex2 KO-2 (11 cells), Tex2 KO-2 resuced by Halo-Tex2 (9 cells), and Tex2 KO-2 rescued by Halo-Tex2-ΔSMP (10 cells) in more than 3 independent assays. Ordinary one-way ANOVA with Tukey’s multiple comparisons test. Mean ± SD. **F.** Representative images of control or two HeLa Tex2-KO clones expressing mApple-Lamp1-phLuorin treated with BafA1 (400nM; 2h) with insets. **G.** Percentage of phLuorin-positive lysosomes in control (10 cells), Tex2 KO-1 (10 cells), Tex2 KO-2 (10 cells) upon BafA1 treatments in 3 independent assays. Ordinary one-way ANOVA with Tukey’s multiple comparisons test. Mean ± SD. Scale bar, 10μm in the whole cell images and 2μm in the insets in (B, C, & F).

The lysosomal defects was specific to Tex2 depletion, since expression of WT Tex2 could completely rescue the phenotype in these two Tex2-KO clones (Fig. 8C, E). Importantly, the lipid transfer-deficient mutant (Tex2-ΔSMP) that was still able to localize to the MCSs failed to rescue the lysosomal defects in two Tex2 KO clones (Fig. 8C, E). The striking difference between WT and lipid transfer-deficient mutant Tex2 in rescue experiments were not due to their expression level, as immunoblot assays showing a similar level of these two proteins (Fig. 8D). These results suggest that Tex2 is required for maintainance of the lysosomal digestive function in a lipid transfer-dependent manner.

Notably, BafA1 treatment did not caused an additive increase in the number of phLuorin-postive LE/lys in Tex2-KO cells (Fig. 8F, G), suggesting that Tex2-KO affects lysosomal functions by hampering the lysosomal pH. In addition, we examined the level of Cathepsin D, a lysosomal aspartyl protease in lysosmal lumen, and we found that the matured form of Cathepsin D was not substantially reduced in Tex2 KO, compared to control (Fig. S8A), suggesting that Tex2 may not directly regulate the sorting or activities of lysosomal hydrolases.

To gain more insights of Tex2 on the digestive functions of lysosomes, we explored the effects of Tex2 KO on autophagy, a fundamental process closely linked to lysosomal digestive capacity (Dikic and Elazar, 2018). We assessed the basal autophagic flow by a specific sensor RFP-LC3-GFP (Fig. 9A) (Kaizuka et al., 2016). Contrary to the control, in which few autophagosomes labeled by GFP-LC3 puncta were found under normal conditions, the number of autophagosomes were markedly increased in two Tex2 KO clones (Figs. 9B, E). Importantly, the accumulation of autophagosomes in Tex2 KO could be almost completely rescued by WT-Tex2 other than Tex2-ΔSMP (Fig. 9C, D, & E). Notably, we found that the increase in autophagosome number resulted from Tex2 KO is likely due to the impaired digestive capacity of lysosomes instead of defective autophagosome-lysosome fusion because we observed that majority of GFP-LC3-labeled autophaosomes were colocalized with Lamp1-labeled LE/lys under either normal or BafA1-treated Tex2-KO cells (Fig. S8B, C). In addition, we confirmed that the flow of Rapamycin-induced autophagy was also blocked at autolysosome stage in the two Tex2-KO clones, as revealed by a strong accumulation of autolysosomes labled by GFP-LC3 and Lysotracker (Fig. 9F, H), which was substantially rescued by WT-Tex2 other than Tex2-ΔSMP (Fig. 9G, H), indicating that the lipid transfer actrivity of Tex2 is required for the proper autophagic flow.

**Fig.9.**
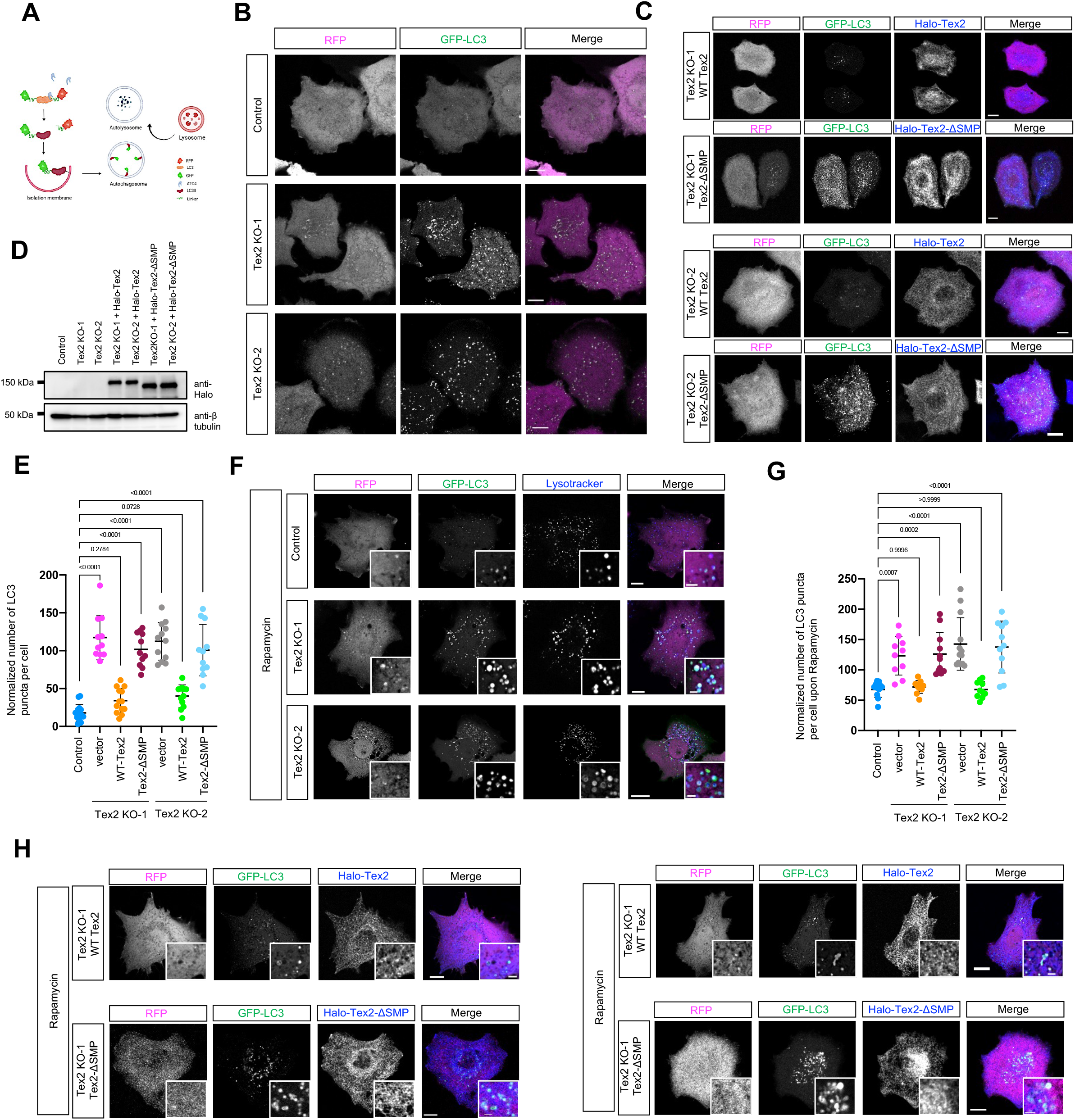
Tex2 depletion results in impaired autophagic flow. **A.** Schematic cartoon of GFP-LC3-RFP as an autophagic flow sensor. **B.** Representative images of control or two HeLa Tex2-KO clones (middle) expressing GFP-LC3-RFP (GFP-LC3 in green; internal control RFP in magenta). **C.** Representative images of two HeLa Tex2-KO clones expressing either Halo-Tex2 (blue) or Halo-Tex2-ΔSMP (blue) and GFP-LC3-RFP with insets. **D.** Western blots demonstrate the level of Halo-Tex2 and Halo-Tex2-ΔSMP in rescue experiments in (**C**). **E.** Normalized number of LC3-positive autophagosomes per cell under normal condition in control (14 cells), Tex2 KO-1 (11 cells), Tex2 KO-1 rescued by Halo-Tex2 (13 cells), Tex2 KO-1 rescued by Halo-Tex2-ΔSMP (11 cells), Tex2 KO-2 (12 cells), Tex2 KO-2 rescued by Halo-Tex2 (12 cells), and Tex2 KO-2 rescued by Halo-Tex2-ΔSMP (11 cells) in 3 independent assays. Ordinary one-way ANOVA with Tukey’s multiple comparisons test. Mean ± SD. **F.** Representative images of control or two HeLa Tex2-KO clones expressing GFP-LC3-RFP under rapamycin treatments (400 nM; 2h) with insets. **G.** Representative images of two HeLa Tex2-KO clones expressing either Halo-Tex2 (blue) or Halo-Tex2-ΔSMP (blue) and GFP-LC3-RFP under rapamycin treatments (400 nM; 2h) with insets. **H.** Normalized number of LC3-positive autophagosomes per cell under rapamycin stimulation in control (11 cells), Tex2 KO-1 (10 cells), Tex2 KO-1 rescued by Halo-Tex2 (10 cells), Tex2 KO-1 rescued by Halo-Tex2-ΔSMP (11 cells), Tex2 KO-2 (13 cells), Tex2 KO-2 rescued by Halo-Tex2 (10 cells), and Tex2 KO-2 rescued by Halo-Tex2-ΔSMP (11 cells) in 3 independent assays. Ordinary one-way ANOVA with Tukey’s multiple comparisons test. Mean ± SD. Scale bar, 10μm in the whole cell images and 2μm in the insets in (B, C, F, & G).

It is intriguing that lysosomal digestive functions are dependent on the lipid transfer activity of Tex2, prompting us to explore whether Tex2 is required for the maintainence of the lipid composition of LE/lys membranes by live-cell microscopy. To begin with, we found that the distribution of PS was not substantially affected by Tex2 KO (Fig. S9A, D), as revealed by GFP-Lact-C2 (Yeung et al., 2008), a PS sensor that specifically marks the PS on the cytoplasmic face of intracellular membranes. Similarly, Tex2 KO did not caused a substantial change in the distribution of either PI(4,5)P_2_ or PI4P on the LE/lys membranes (Fig. S9B-D), as shown by the PI(4,5)P_2_ sensor GFP-PLC8-PH (Stauffer et al., 1998), or the PI4P sensor GFP-OSBP-PH (Levine and Munro, 2002), respectively.

PI3P is primarily found in microdomains of early endosomes, and plays essential roles in endosomal maturation and functions (Gillooly et al., 2003; Marat and Haucke, 2016). Supporting this notion, we found that, in control cells, p40PX-GFP, a PI3P sensor (Kanai et al., 2001), primarily decorated the membranes of early endosomes, which was marked by early endosome antigen 1 (EEA1) (pearson’s correlation coefficient= 0.86) (Fig. S9E & F). In contrast, we found that PI3P was significantly detected on the membranes of LE/lys in two Tex2-KO clones, as revealed by the co-localization between the PI3P sensor p40PX-GFP and the LE/lys marker Lamp1-mCh (Fig. 10A, D). Importantly, the defect could be significantly restored by introduction of WT Tex2 to the two Tex2-KO cells, while the expression of Tex2-ΔSMP had a negligible effect in rescue experiments (Fig. 10B, C, D), suggesting that lipid transfer activities of Tex2 is required endosomal maturation.

**Fig.10.**
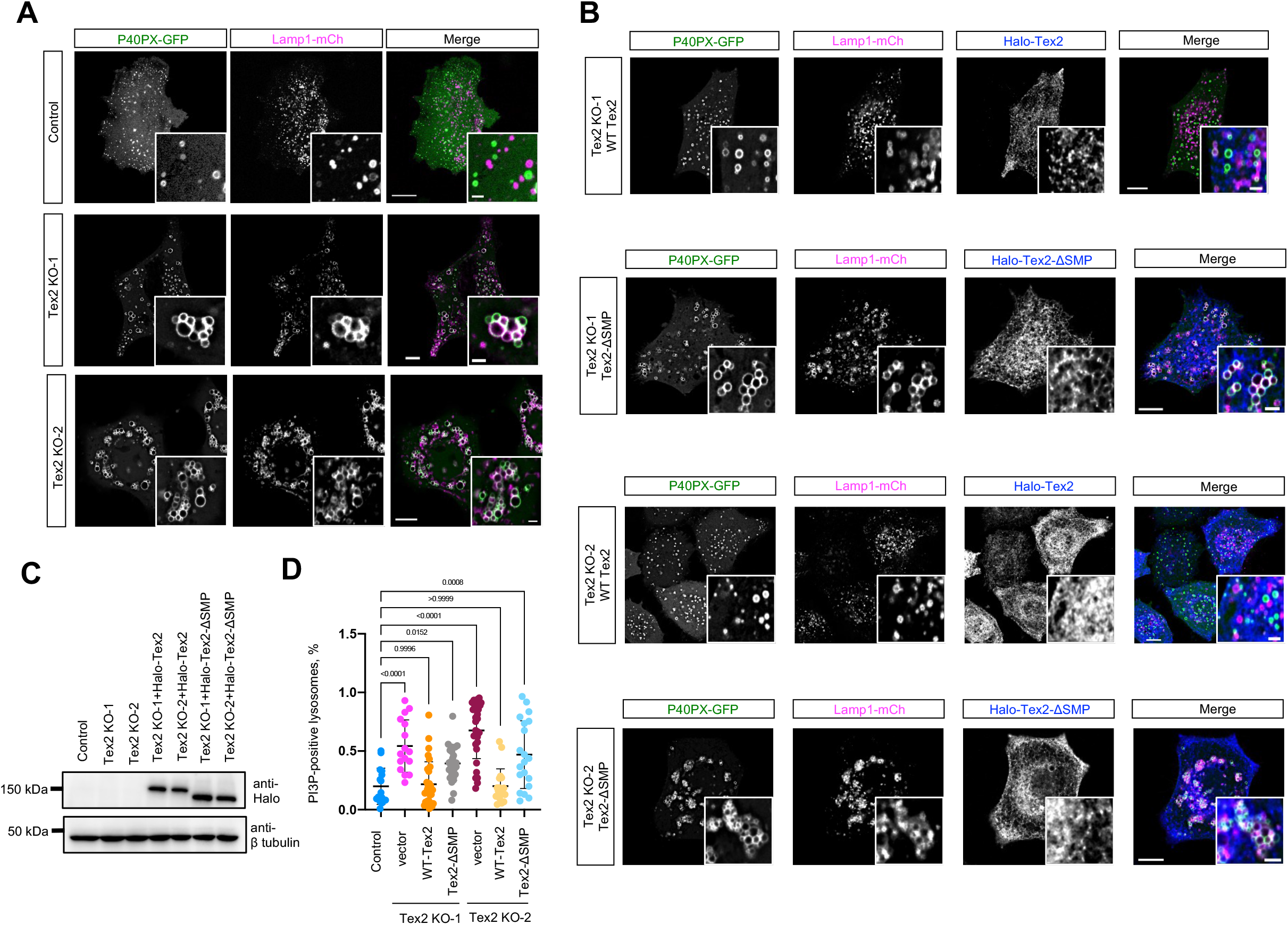
Tex2 depletion increases PI3P on LE/lys membranes. **A.** Representative images of control or two HeLa Tex2-KO clones expressing PI3P sensor P40PX-GFP (green) and Lamp1-mCh (magenta) with insets. **B.** Representative images of two HeLa Tex2-KO clones expressing either Halo-Tex2 (blue) or Halo-Tex2-ΔSMP (blue) along with P40PX-GFP (green) and Lamp1-mCh (magenta) with insets. **C.** Western blots demonstrate the level of Halo-Tex2 and Halo-Tex2-ΔSMP in rescue experiments. **D.** Number of P40PX-positive lysosomes per cell in control (16 cells), Tex2 KO-1 (16 cells), Tex2 KO-1 rescued by Halo-Tex2 (30 cells), Tex2 KO-1 rescued by Halo-Tex2-ΔSMP (27 cells), Tex2 KO-2 (30 cells), Tex2 KO-2 rescued by Halo-Tex2 (18 cells), and Tex2 KO-2 rescued by Halo-Tex2-ΔSMP (20 cells) in 3 independent assays. Ordinary one-way ANOVA with Tukey’s multiple comparisons test. Mean ± SD. Scale bar, 10μm in the whole cell images and 2μm in the insets in (A, B).

## Discussion

In this manuscript, we propose a model in which a LTP Tex2, localizing exclusively to tubular ER, is recruited to ER-LE/lys MCSs via LE/lys-resident PI phosphatases TMEM55, and the recruitment of Tex2 by TMEM55 is regulated by PI4KII activities. The loss of lipid transfer activity of Tex2 at these contacts results in severe defects in lysosomal digestive functions, impaired autophagical flow, as well as an aberrant accumulation of PI3P on the LE/lys membranes (Fig. 11).

**Fig. 11.**
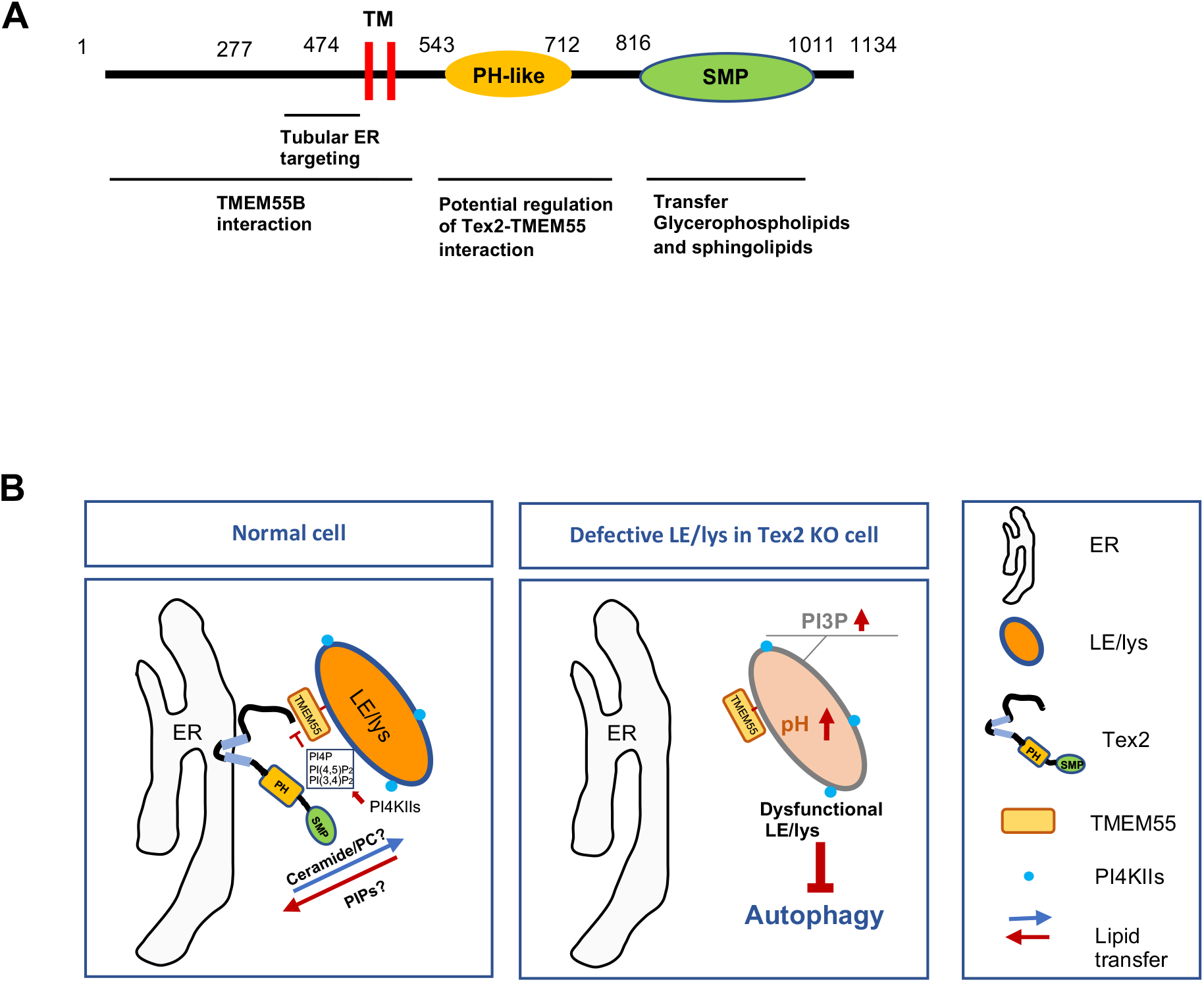
Working model of Tex2 functions at ER-LE/lys MCSs. **A.** Functional dissection of Tex2. Tex2-NT (1-517) containing essential residues 1-277 is required for binding the catalytic motif CX_5_R of TMEM55. The residues 278-514 is required for targeting Tex2 to tubular ER. The SMP domain is likely responsible for transferring glycerophospholipids and sphingolipids between the ER and LE/lys. The PH domain of Tex2 may be involved in the regulation of Tex2-TMEM55 interactions. **B.** Tex2 resides exclusively on tubular ER, is recruited to ER-LE/lys MCSs via LE/lys-resident PI phosphatases TMEM55. The substrate of TMEM55, PI(4,5)P2, promotes Tex2-TMEM55B interactions. The loss of lipid transfer activity of Tex2 at these contacts resulted in severe defects in lysosomal digestive functions, impaired autophagic flow, and an aberrant accumulation of PI3P on LE/lys membranes.

ER tubules has evolved to play specific and fundamental roles in mammalian cells. One of such functions is to synthesize majority of lipids and transfer them to other organelles for organelle biogenesis, trafficking and alleviation of lipitoxicity (Prinz et al., 2020). Our findings coordiante ER tubule formation, endosome maturation, PIPs on LE/lys membranes, and lipid transfer across ER-LE/lys MCSs.

To begin with, our findings demonstrate the localization of Tex2 on tubular ER and the underlying mechanism. The RTN proteins localize exclusively to tubular ER via a conserved C-terminal reticulon homology domain (RHD), which forms a characteristic hairpin TM domain that targets RTN proteins to regions of high membrane curvature including tubular ER membranes (Voeltz et al., 2006). We found that Tex2 is exclusively localized to tubular ER via a region (residues 277-474) of the NT adjacent to TM domain. The question of how the region targets Tex2 to ER tubules remains elusive. We found that two potential amphipathic helices (APH) in this region, but disruption of these two APH by point mutations did not substantially reduce the targeting of Tex2 to ER tubules (unpublished results). Notably, overexpression of Tex2 could counteract the effects of Climp63 in the expansion of ER sheets at cell periphery. However, Tex2 KO appears to not substantially affect the ER tubule-sheet ratio at cell pheriphery. It may suggest that Tex2 is not required for ER tubule formation in cells, which is different form the role of RTN4, the prototype of RTN proteins that generates and maintains the structure of the tubular ER network (Voeltz et al., 2006). The specific localization of Tex2 to tubular ER extends the functional significance of ER tubules in LTP-mediated lipid transport at ER-associated MCSs.

Intracellular LTPs mediate lipid transport between opposing membranes at MCSs through two modes of action – shuttling or bridging. Shuttle transporters typically extract one or two lipid molecules from the membrane of the donor organelle, solubilize it during transport through the cytosol and deposit it in the acceptor organelle membrane, while bridge transporters feature an extended channel, most likely lined with hydrophobic residues that bind tens of lipids at once (Li et al., 2020; Ugur et al., 2020; Wong et al., 2019). In addition to Tex2, a complex of multiple SMP domains are suggested to act as a shuttle transporter for glycerophospholipids and/or ceramides across MCSs in yeast and metazoen, including maintenance of mitochondrial morphology protein 1 (Mmm1) of ERMES (Kornmann et al., 2009) and Nvj2 (Liu et al., 2017)in yeast, and E-Syts (Bian et al., 2018; Giordano et al., 2013), Transmembrane protein 24 (TMEM24; also known as C2CD2L) (Lees et al., 2017), and PDZD8 (Gao et al., 2022). To ensure a productivity of lipid shuttling across MCSs, a directional lipid transfer should be fullfilled. Nvj2p, the yeast homolog of Tex2, may mediates a directional ceramide transfer from the ER to the Golgi apparatus for sphingolipid synthesis during ER stress, possibly driven by ceramide gradients (Liu et al., 2017). The direction and driving force of Tex2-mediated lipid transport is unknown. It is plausible that the PI4,5P_2_ to PI5P conversion catalyzed by TMEM55B may be coupled to the lipid transfer of Tex2, and contributes to a directional lipid transport at ER-LE/lys MCSs in a similar manner as the SMP-containing LTP TMEM24, which directionally transport phosphatidylinositols over other phospholipids at ER-PM MCSs (Lees et al., 2017). Supporting this hypothesis, our results showed that PI3P was aberrantly accumulated on the membranes of LE/lys (Fig. 10), and we found that the SMP domain of Tex2 might bind diverse PIPs, including PI3P, PI4P, PI5P, PI(3,5)P_2_, PI(4,5)P_2_, and PI(3,4,5)P_3_ (unpublished data). Whether Tex2 binds these PIPs and other phospholipids (PC or Ceramide) via same sites of SMP? Whether Tex2 transfers PIPs, for example PI5P, over PC or Ceramide in cells? These critical questions warrant fututre investigations.

Tex2 is highly expressed in the testis, and our findings highlighting the importance of lipid transfer activities of Tex2 in regulating lysosomal digestive function raise questions of whether Tex2 is required in the biogenesis of acrosome, a specialized lysosome-like organelles containing hydrolases in sperm for egg penetration (Moreno and Alvarado, 2006), warranting further investigations.

## Method

### Cell culture, transfection, RNAi

The African green monkey kidney fibroblast-like COS7 cell line (ATCC), human cervical cancer HeLa cells (ATCC), and human embryonic kidney 293T (ATCC) were grown in DMEM (Invitrogen) supplemented with 10% fetal bovine serum (Gibco). All of the cell lines used in this study were confirmed free of mycoplasma contamination.

Transfection of plasmids and RNAi oligos was carried out with Lipofectamine 2000 and RNAi MAX, respectively. For transfection, cells were seeded at 4 x 10^5^ cells per well in a six-well dish ∼16 h before transfection. Plasmid transfections were performed in OPTI-MEM (Invitrogen) with 2 μl Lipofectamine 2000 per well for 6 h, followed by trypsinization and replating onto glass-bottom confocal dishes at ∼3.5 x 10^5^ cells per well. Cells were imaged in live-cell medium (DMEM with 10% FBS and 20 mM Hepes with no penicillin or streptomycin) ∼16–24 h after transfection. For all transfection experiments in this study, the following amounts of DNA were used per 3.5 cm well (individually or combined for co-transfection): 1500 ng for GFP-Tex2 and its mutants, 500 ng for Halo-TMEM55A/B and its mutants, Lamp1-Halo; 1500 ng for ER-tagRFP. For siRNA transfections, cells were plated on 3.5 cm dishes at 30–40% density, and 2 μl Lipofectamine RNAimax (Invitrogen) and 50 ng siRNA were used per well. At 48 h after transfection, a second round of transfection was performed with 50 ng siRNAs. Cells were analyzed 24 h after the second transfection for suppression.

### Plasmid

Tex2 (NM_018469.5), TMEM55A (NM_018710.3), TMEM55B (NM_001100814.3), E-syt1, ALT1, ARL6IP1, E-Syt1, RTN4, Reep5 were cloned from HeLa cDNA. The ORFs of Tex2 and TMEM55B, and E-syt1 were cloned into mEGFP-C1/Halo-C1 vector between the BglII and SacI. The ORFs of Reep5 and RTN4 were cloned between NheI and SacI of Halo-N1. GST-TMEM55B was cloned into PGEX-2T vector between the BamHI and EcoRI. 14xHis-NEDD8-Tex2-SMP was cloned into 14xHis-NEDD8 vector between the BamHI and HindIII. ER-tagRFP and Lamp1-mCh were previously described (Gao et al., 2022; Ji et al., 2017). p40PX-EGFP (addgene 19010), GFP-Lact-C2 (addgene 22852), CFP-C1-PLC*δ*-PH (addgene 21262), pBGPa-CMV-GFP-OSBP-PH (addgene 58724), mApple-Lamp1-phLuorin-N-8 (addgene 54918), and PMRX-IP-GFP-LC3-RFP (addgene 84573) were purchased from Addgene.

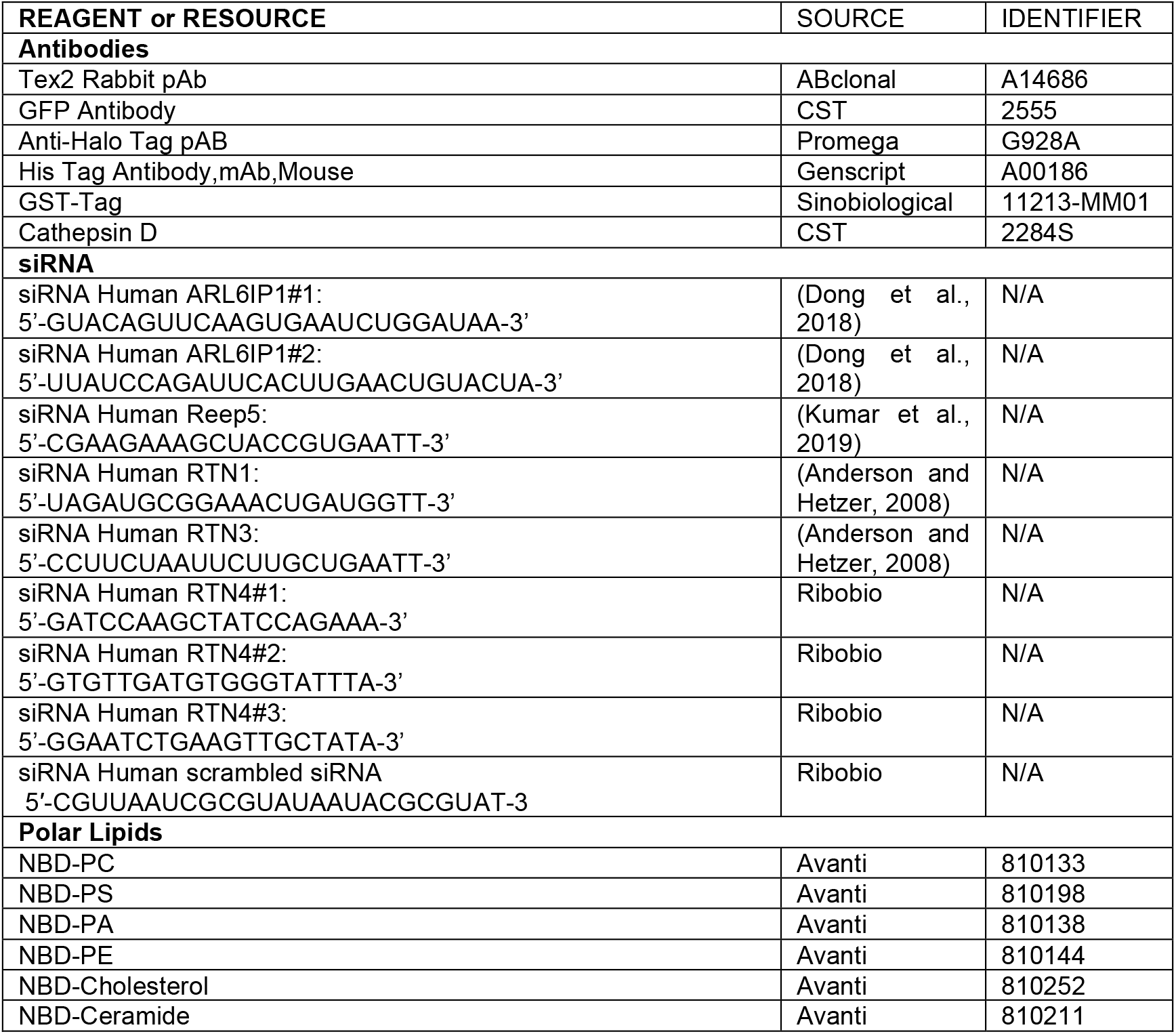
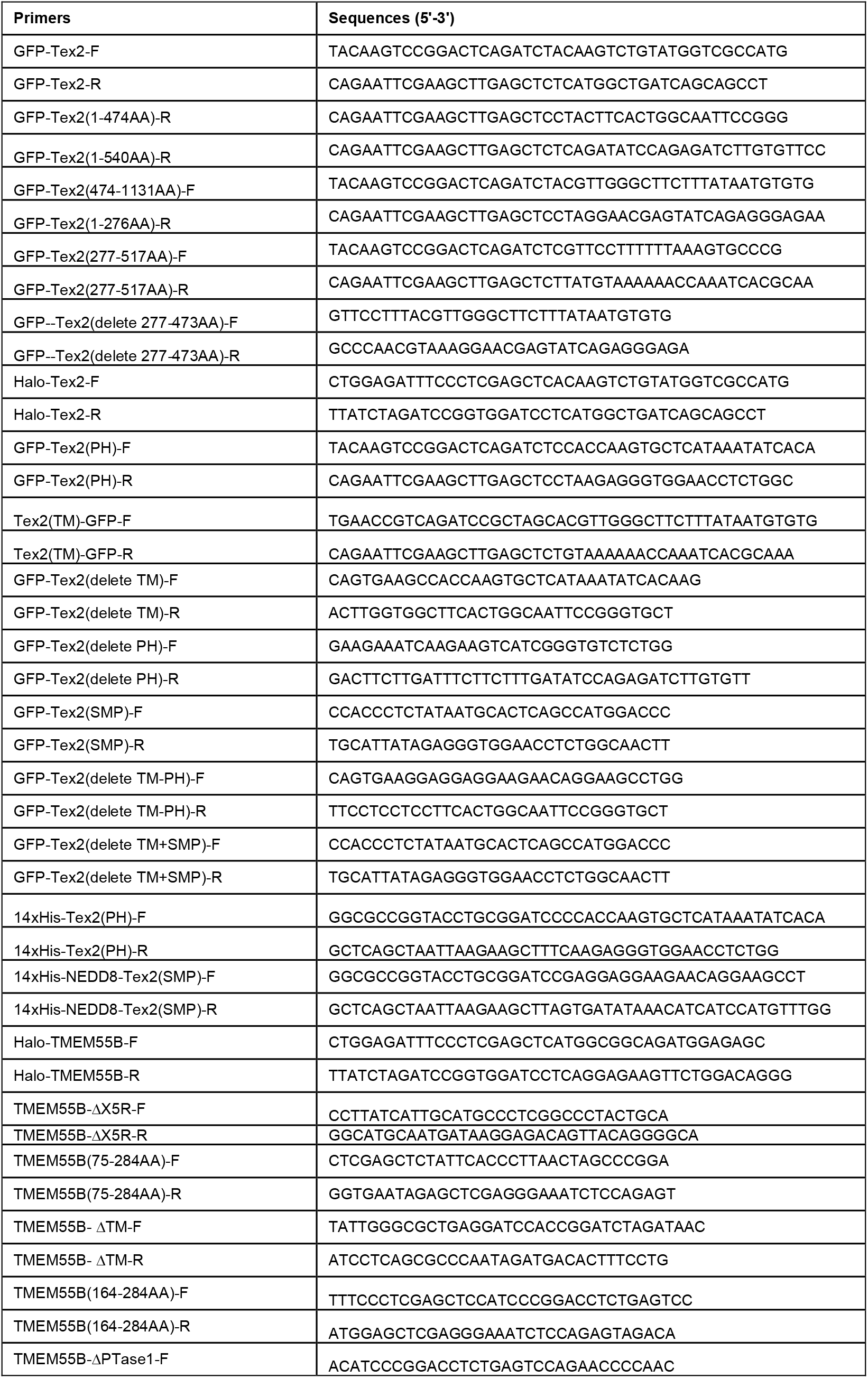
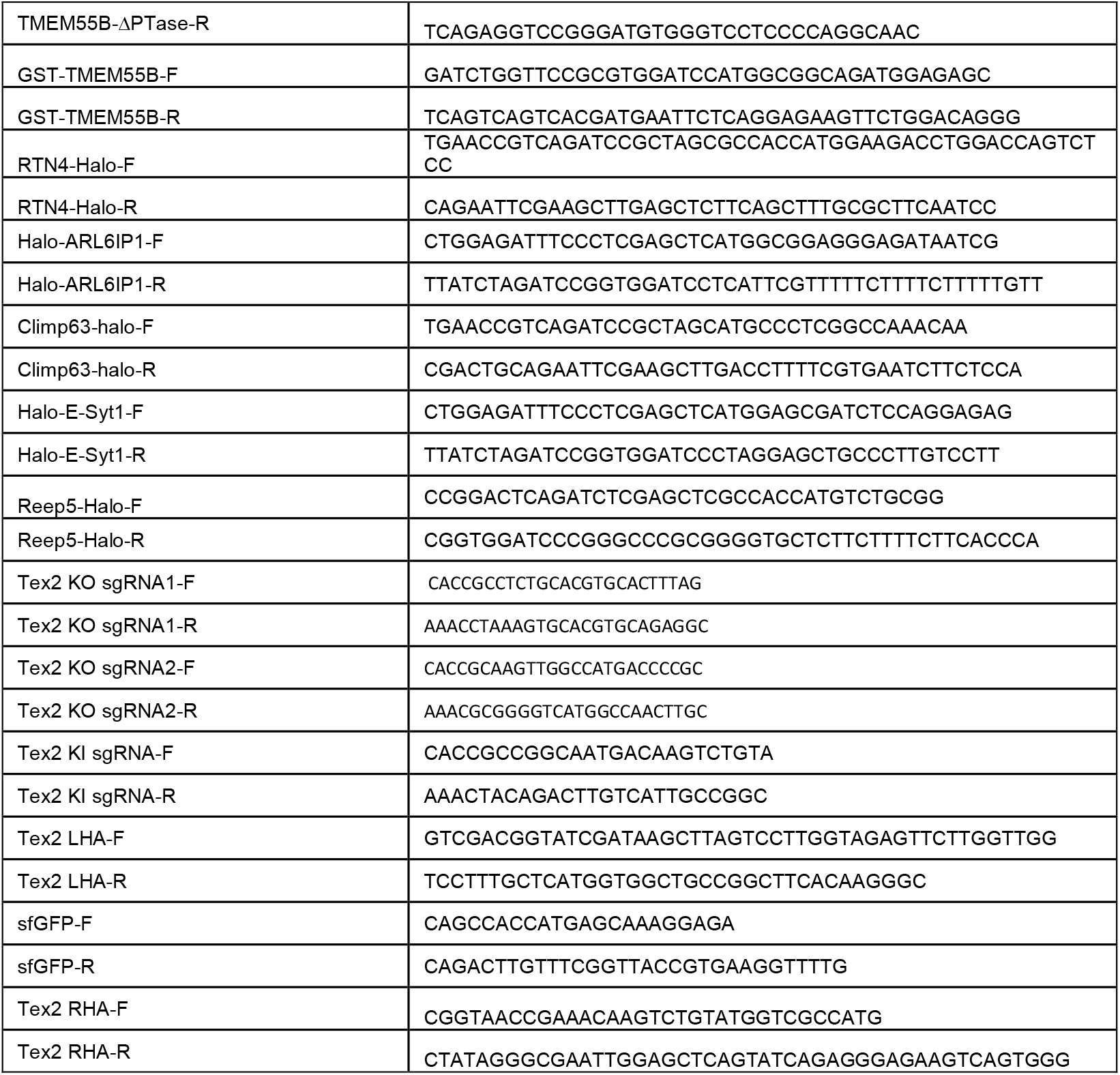

### GFP-trap assay

GFP trap (GTA-100, ChromoTek) was used for detection of protein–protein interactions and the GFP-Trap assays were performed according to the manufacturer’s protocol. 5% input was used in GFP traps unless otherwise indicated.

### GST Tag and His Tag protein purification

GST and His constructs were transformed into Escherichia coli BL21 (DE3) cells, and cells were incubated at 37°C until the optical density (OD) at 600 nm reached 0.6–0.8. Subsequently, cells were incubated at 16°C for another hour, followed by induction with 1 mM IPTG overnight at 16°C.Cells were lysed via sonication. GST fusion proteins were purified via the GST-tag Protein Purification kit (C600031-0025, Sangon, China), His fusion proteins were purified via the Ni-NTA Sefinose (TM) Resin Purification kit (G600033-0100, Sangon, China)

### In vitro lipid-binding assay

1 μl of either NBD-labeled PS, PE, PC, PA, Cholesterol, or Ceramide (1 mg/ml in methanol) was incubated with 19 μl purified 14xHis-Tex2-SMP (1 mg/ml) for 2 h at 4°C. Samples were visualized on 10% native PAGE gels. NBD fluorescence was visualized using Bio-Rad ChemiDoc^TM^ MP, and comigrated proteins were visualized by Coomassie blue staining.

### In vitro Pull-down assays of GFP-Tex2 and GST-TMEM55B

HEK293 cells transiently transfected with GFP-Tex2 were lysed in high-salt lysis buffer (RIPA buffer containing 500 mM NaCl, proteasome inhibitors and PMSF). GFP-Trap beads were used to pellet GFP-Tex2 from cell lysates, followed by washing with high-salt lysis buffer for 10 times. The GFP-Tex2 beads were incubated with different amounts of Ceramide or PS (0, 0.2, 0.4, 0.6, 0.8, 1.0 mg) for 2 h at 4°C, and then were incubated with Purified GST–TMEM55B or GST-only overnight at 4°C, respectively, followed by washing beads with freshly prepared HNM buffer (20 mM Hepes, pH 7.4, 0.1 M NaCl, 5 mM MgCl2, 1 mM DTT and 0.2% NP-40). GFP-Tex2 beads were resuspended in 100 μL 2 x SDS-sampling buffer. Re-suspended beads were boiled for 10 min at 95°C to dissociate protein complexes from beads. Western blotting was performed using anti-GFP, GST or TMEM55B antibodies. The Coomassie staining was performed for purified GST-TMEM55B.

### PIP Strip assays

The PIP Strips (P-6001) were blocked by TBS-T + 3% fatty acid–free BSA, and then were gently agitated for 1 h at room temperature, followed by an incubation with purified His-Tex2-PH (0.5 µg/mL) in TBS-T + 3% fatty acid–free BSA overnight at 4°C. After washing the PIP Strips with TBS-T + 3%fatty acid–free BSA three times under gentle agitation for 10 min each time, PIP strips were incubated with the anti-His antibodies overnight at 4°C, followed by repeated washing steps.

### Live imaging by high-resolution confocal microscopy

Cells were grown on 35 mm glass-bottom confocal MatTek dishes, and the dishes were loaded to a laser scanning confocal microscope (LSM900, Zeiss, Germany) equipped with multiple excitation lasers (405 nm, 458 nm, 488 nm, 514 nm, 561 nm and 633 nm) and a spectral fluorescence GaAsP array detector. Cells were imaged with the 63×1.4 NA iPlan-Apochromat 63 x oil objective using the 405 nm laser for BFP, 488 nm for GFP, 561nm for OFP, tagRFP or mCherry and 633nm for Janilia Fluo® 646 HaloTag® Ligand.

### Mass spectrometry for identification of GFP-Tex2-interacting proteins

The identification of GFP-Tex2-interacting proteins by MS was described in our previous study (Gao et al., 2022). Briefly, the bound proteins were extracted from GFP-Trap agarose beads using SDT lysis buffer (4% SDS, 100 mM DTT, 100 mM Tris-HCl pH 8.0), followed by sample boiling for 3 min and further ultrasonicated. Undissolved beads were removed by centrifugation at 16,000 g for 15 min. The supernatant, containing proteins, were collected. Protein digestion was performed with FASP method. Briefly, the detergent, DTT and IAA in UA buffer was added to block-reduced cysteine. Finally, the protein suspension was digested with 2 µg trypsin (Promega) overnight at 37°C. The peptide was collected by centrifugation at 16,000 g for 15 min. The peptide was desalted with C18 StageTip for further LC-MS analysis. LC-MS/MS experiments were performed on a Q Exactive Plus mass spectrometer that was coupled to an Easy nLC (Thermo Fisher Scientific). Peptide was first loaded to a trap column (100 µm x 20 mm, 5 µm, C18, Dr Maisch GmbH, Ammerbuch, Germany) in buffer A (0.1% formic acid in water). Reverse-phase high-performance liquid chromatography (RP-HPLC) separation was performed using a self-packed column (75 µm x 150 mm; 3 µm ReproSil-Pur C18 beads, 120 Å, Dr Maisch GmbH, Ammerbuch, Germany) at a flow rate of 300 nl/min. The RP-HPLC mobile phase A was 0.1% formic acid in water, and B was 0.1% formic acid in 95% acetonitrile. The gradient was set as following: 2%–4% buffer B from 0 min to 2 min, 4% to 30% buffer B from 2 min to 47 min, 30% to 45% buffer B from 47 min to 52 min, 45% to 90% buffer B from 52 min and to 54 min, and 90% buffer B kept until to 60 min. MS data was acquired using a data-dependent top20 method dynamically choosing the most abundant precursor ions from the survey scan (350– 1800 m/z) for HCD fragmentation. A lock mass of 445.120025 Da was used as internal standard for mass calibration. The full MS scans were acquired at a resolution of 70,000 at m/z 200, and 17,500 at m/z 200 for MS/MS scan. The maximum injection time was set to 50 ms for MS and 50 ms for MS/ MS. Normalized collision energy was 27 and the isolation window was set to 1.6 Th. Dynamic exclusion duration was 60 s. The MS data were analyzed using MaxQuant software version 1.6.1.0. MS data were searched against the UniProtKB Rattus norvegicus database (36,080 total entries, downloaded 08/14/2018). Trypsin was seleted as the digestion enzyme. A maximum of two missed cleavage sites and the mass tolerance of 4.5 ppm for precursor ions and 20 ppm for fragment ions were defined for database search. Carbamidomethylation of cysteines was defined as a fixed modification, while acetylation of protein N-terminal, oxidation of Methionine were set as variable modifications for database searching. The database search results were filtered and exported with a <1% false discovery rate (FDR) at peptide-spectrum-matched level, and protein level, respectively.

### Mass spectrometry for identification of GFP-Tex2-associated lipids in HEK293 cells

The identification of GFP-Tex2-assocaiting lipids in HEK293 cells was described in our previous study (Gao et al., 2022). Briefly, To extract lipids, 1 ml methyl tert-butyl ether (MTBE) was added to GFP-Trap agarose beads (Chromoteck) and the samples were shaken for 1 h at room temperature. Next, phase separation was induced by adding 250 µL water, letting it sit for 10 min at room temperature and centrifuging for 15 min at 14,000 g, 4°C. Because of the low density and high hydrophobicity of MTBE, lipids and lipophilic metabolites are mainly extracted to the upper MTBE-rich phase. The lipid was transferred to fresh tubes and dried with nitrogen. Additionally, to ensure data quality for metabolic profiling, quality control (QC) samples were prepared by pooling aliquots from representative samples for all of the analysis samples, and were used for data normalization. QC samples were prepared and analyzed with the same procedure as that for the experiment samples in each batch. Dried extracts were then dissolved in 50% acetonitrile. Each sample was filtered with a disposable 0.22 μm cellulose acetate and transferred into 2 ml HPLC vials and stored at -80°C until analysis. For UHPLC-MS/MS analysis, lipid analysis was performed on Q Exactive orbitrap mass spectrometer (Thermo Fisher Scientific) coupled to a UHPLC system Ultimate 3000 (Thermo Fisher Scientific). Samples were separated using a Hypersil GOLD C18 column (100 x 2.1 mm, 1.9 µm) (Thermo Fisher Scientific). Mobile phase A was prepared by dissolving 0.77 g of ammonium acetate to 400 ml of HPLC-grade water, followed by adding 600 ml of HPLC-grade acetonitrile. Mobile phase B was prepared by mixing 100 ml of acetonitrile with 900 ml isopropanol. The flow rate was set as 0.3 mL/min. The gradient was 30% B for 0.5 min and was linearly increased to 100% in 10.5 min, and then maintained at 100% in 2 min, and then reduced to 30% in 0.1 min, with 4.5 min re-equilibration period employed. Both electrospray ionization (ESI) positive-mode and negative-mode were applied for MS data acquisition. The positive mode of spray voltage was 3.0 kV and the negative mode 2.5 kV. The ESI source conditions were set as follows: heater temperature of 300°C, Sheath Gas Flow rate, 45arb, Aux Gas Flow Rate, 15 arb, Sweep Gas Flow Rate, 1arb, Capillary Temp, 350°C, S-Lens RF Level, 50%. The full MS scans were acquired at a resolution of 70,000 at m/z 200, and 17,500 at m/z 200 for MS/MS scans. The maximum injection time was set to for 50 ms for MS and 50 ms for MS/MS. MS data was acquired using a data-dependent Top10 method dynamically choosing the most abundant precursor ions from the survey scan (200–1500m/z) for HCD fragmentation. Stepped normalized collision energy was set as 15, 25, 35 and the isolation window was set to 1.6 Th. QC samples were prepared by pooling aliquots that were representative of all samples under analysis, and used for data normalization. Blank samples (75% acetonitrile in water) and QC samples were injected every six samples during acquisition.

For data preprocessing and filtering, lipids were identified and quantified using LipidSearch 4.1.30 (Thermo, CA). Mass tolerance of 5 ppm and 10 ppm were applied for precursor and product ions. Retention time shift of 0.25 min was performed in ‘alignment’. M-score and chromatographic areas were used to reduce false positives. The lipids with less than 30% relative standard deviation (RSD) of MS peak area in the QC samples were kept for further data analysis. SIMCAP software (Version 14.0, Umetrics, Sweden) was used for all multivariate data analyses and modeling. Data were mean-centered using Pareto scaling. Models were built on principal component analysis (PCA), orthogonal partial least-square discriminant analysis (PLS-DA) and partial least-square discriminant analysis (OPLS-DA). All the models evaluated were tested for over fitting with methods of permutation tests. The descriptive performance of the models was determined by R2X (cumulative) [perfect model: R2X (cum)=1] and R2Y (cumulative) [perfect model: R2Y (cum)=1] values while their prediction performance was measured by Q2 (cumulative) [perfect model: Q2 (cum)=1] and a permutation test (n=200). The permuted model should not be able to predict classes – R2 and Q2 values at the Y-axis intercept must be lower than those of Q2 and the R2 of the non-permuted model. OPLS-DA allowed the determination of discriminating metabolites using the variable importance on projection (VIP). The VIP score value indicates the contribution of a variable to the discrimination between all the classes of samples. Mathematically, these scores are calculated for each variable as a weighted sum of squares of PLS weights. The mean VIP value is 1, and usually VIP values over 1 are considered as significant. A high score is in agreement with a strong discriminatory ability and thus constitutes a criterion for the selection of biomarkers. The discriminating metabolites were obtained using a statistically significant threshold of variable influence on projection (VIP) values obtained from the OPLS-DA model and two-tailed Student’s t-test (P-value) on the normalized raw data at univariate analysis level. The P-value was calculated by one-way analysis of variance (ANOVA) for multiple groups analysis. Metabolites with VIP values greater than 1.0 and P-value less than 0.05 were considered to be statistically significant metabolites. Fold change was calculated as the logarithm of the average mass response (area) ratio between two arbitrary classes. On the other side, the identified differential metabolites were used to perform cluster analyses with R package.

### Statistical analysis

All statistical analyses and p-value determinations were performed in GraphPad Prism6. All the error bars represent Mean ± SD. To determine p-values, ordinary one-way ANOVA with Tukey’s multiple comparisons test were performed among multiple groups and a two-tailed unpaired student t-test was performed between two groups.

## Acknowledgements

We thank Anbing Shi and Yanling Yan for insightful suggestions. We thank Qing Tian and Linfang Yang (Huazhong University of Science and Technology, HUST) and Ping Liu (the Optical Bioimaging Core Facility of WNLO-HUST) for imaging assistance. W.J. was supported by National Natural Science Foundation of China (32122025), and the Program for HUST Academic Frontier Youth Team (2018QYTD11).

## Author contributions

Y.D. and W.J. conceived the project and designed the experiments. Y.D. performed the experiments. Y.D. W. C., L.D., and W.J. analyzed and interpreted the data. W.J. prepared the manuscript with inputs and approval from all authors.

## Competing interests

The authors declare no competing interests.

## Data and materials availability

All the data and relevant materials, including reagents and primers, that supports the findings of this study are available from the corresponding author upon reasonable request.

---

**Fig. S1.**
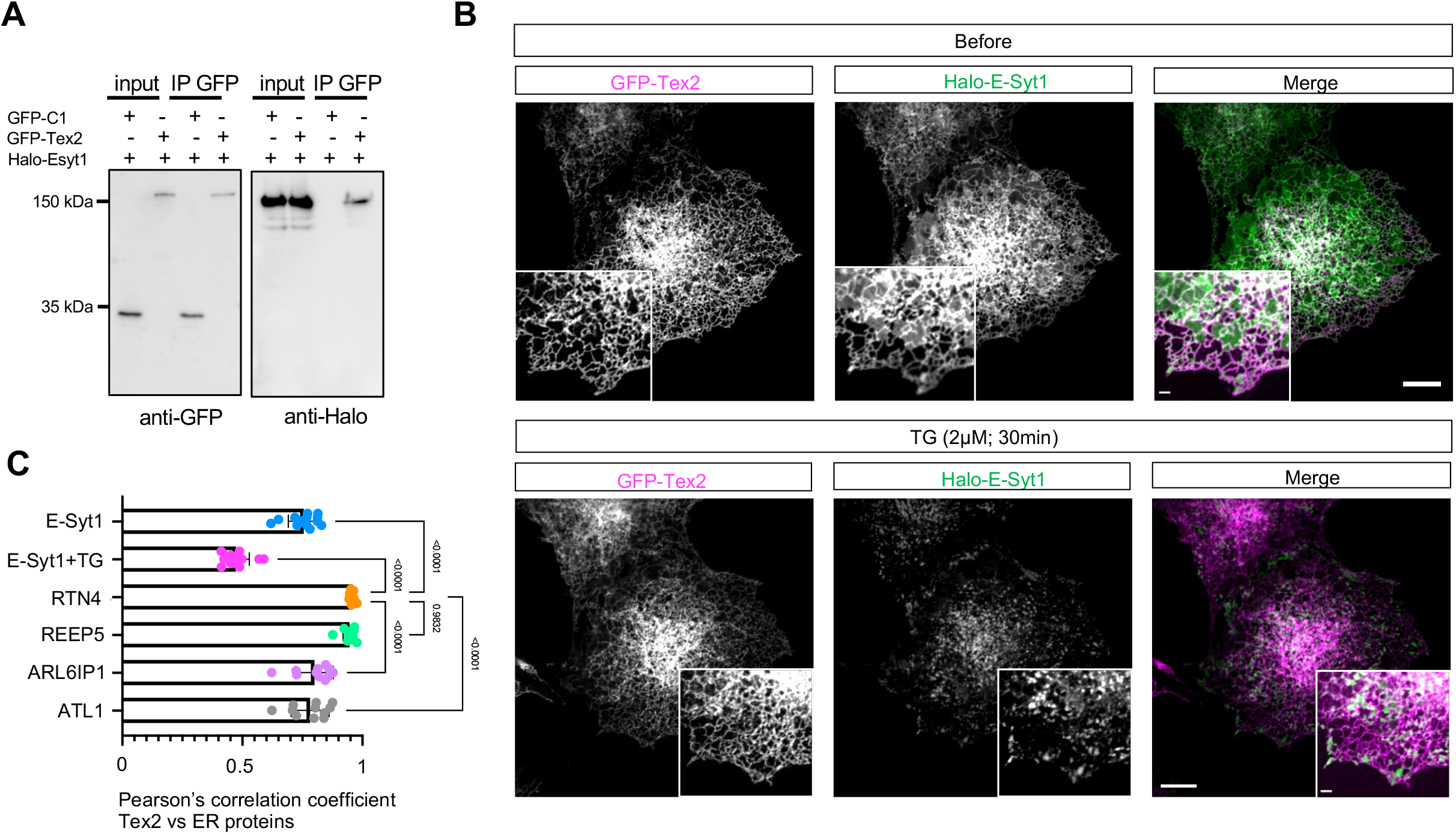
The interaction between Tex2 and E-Syt1. **A.** GFP-Trap assays demonstrate an interaction between GFP-Tex2 and Halo-E-Syt1 in COS7 cells. **B.** Representative images of a COS7 cell expressing GFP-Tex2 (magenta) and Halo-E-Syt1 (green) before (top panel) and after (bottom panel) TG treatment (2 μM; 30 min). **C.** Pearson’s correlation coefficient of Tex2 vs ER-resident proteins, including E-Syt1 (13 cells); E-Syt1 upon TG (13 cells); RTN4 (12 cells); Reep5 (12 cells); ARLIP6 (11 cells); and ATL1 (11 cells) in 3 independent experiments. Ordinary one-way ANOVA with Tukey’s multiple comparisons test. Mean ± SD. Scale bar, 10μm in the whole cell images and 2μm in the insets in (B).

**Fig. S2.**
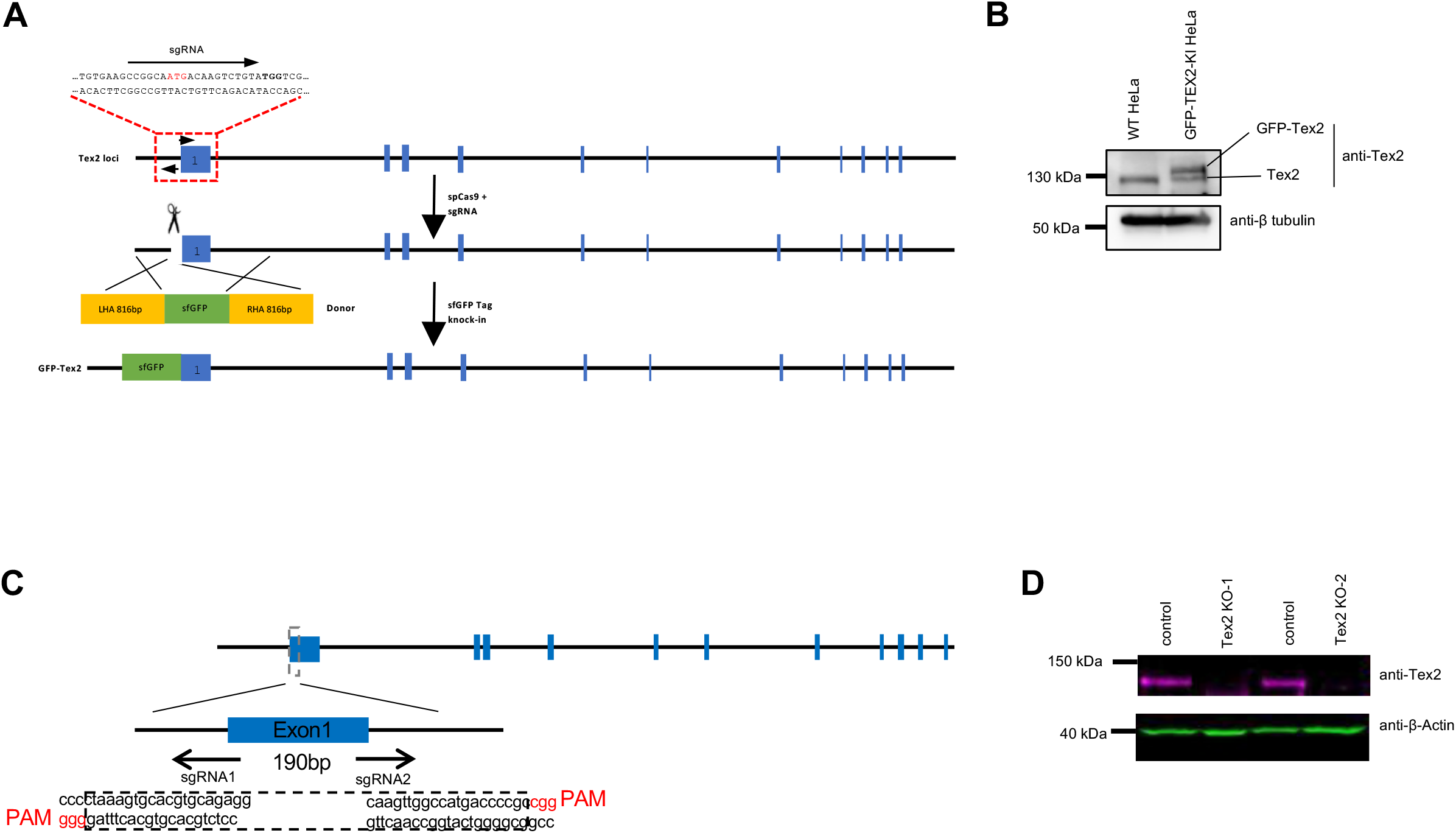
Genome editing of Tex2. A. CRISPR knock-in of GFP (A206K) Tag at N-terminus of Tex2 locus in HeLa cells. The underlined letters indicate the PAM (TGG) for spCas9; ATG in red in sgRNA showing the start codon of Tex2. B. Western blots of GFP-Tex2-KI for single cell clone from (**A**). C. CRISPR knock-out of Tex2 in HeLa cells (Tex2-KO). Two sgRNAs are used with the underlined letters indicate the PAMs (GGG for sgRNA1 and CGG for sgRNA2) for spCas9. D. Western blots of two Tex2-KO clones from (**C**).

**Fig. S3.**
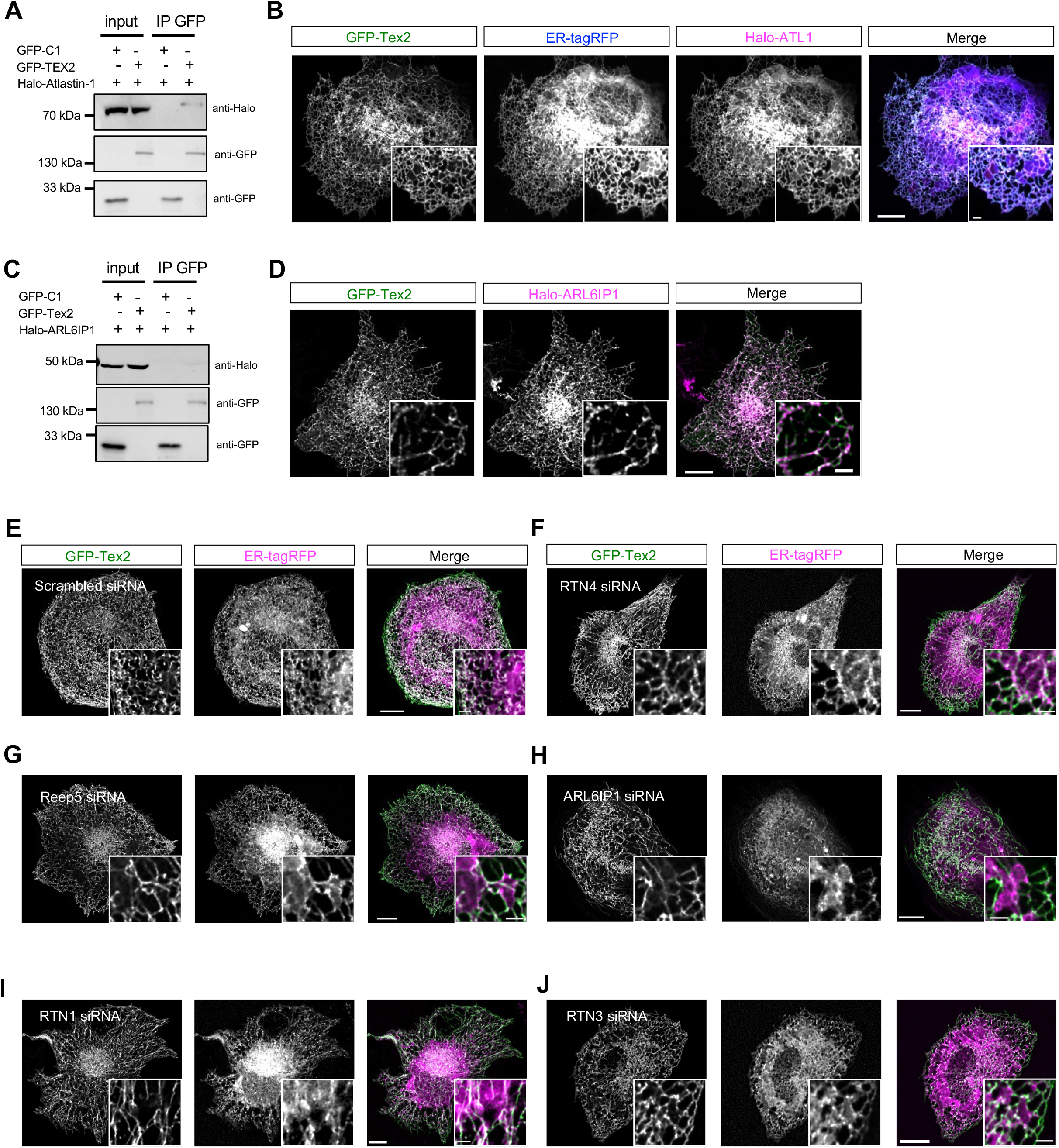
The targeting of Tex2 to tubular ER appears to be independent of ER tubule-shaping proteins. **A.** GFP-Trap assays demonstrate a weak interaction between GFP-Tex2 and Halo-ATL1 in COS7 cells. **B.** Representative images of a COS7 cell expressing GFP-Tex2 (green), Halo-ATL1 (magenta), and ER-tagRFP (blue) with insets. **C.** GFP-Trap assays demonstrate no strong interaction between GFP-Tex2 and Halo-ARL6IP1 in COS7 cells. **D.** Representative images of a COS7 cell expressing GFP-Tex2 (green), Halo-ARL6IP1 (magenta), and ER-tagRFP (blue) with insets. **E-F**. Representative images of COS7 cells expressing GFP-Tex2 (green), ER-tagRFP (magenta) with insets upon treatments of scrambled (E), RTN4 (F), REEP5 (G), ARL6IP1 (H), RTN1 (I), and RTN3 (J) siRNAs. Scale bar, 10μm in the whole cell images and 2μm in the insets in (B, D, E-J).

**Fig. S4.**
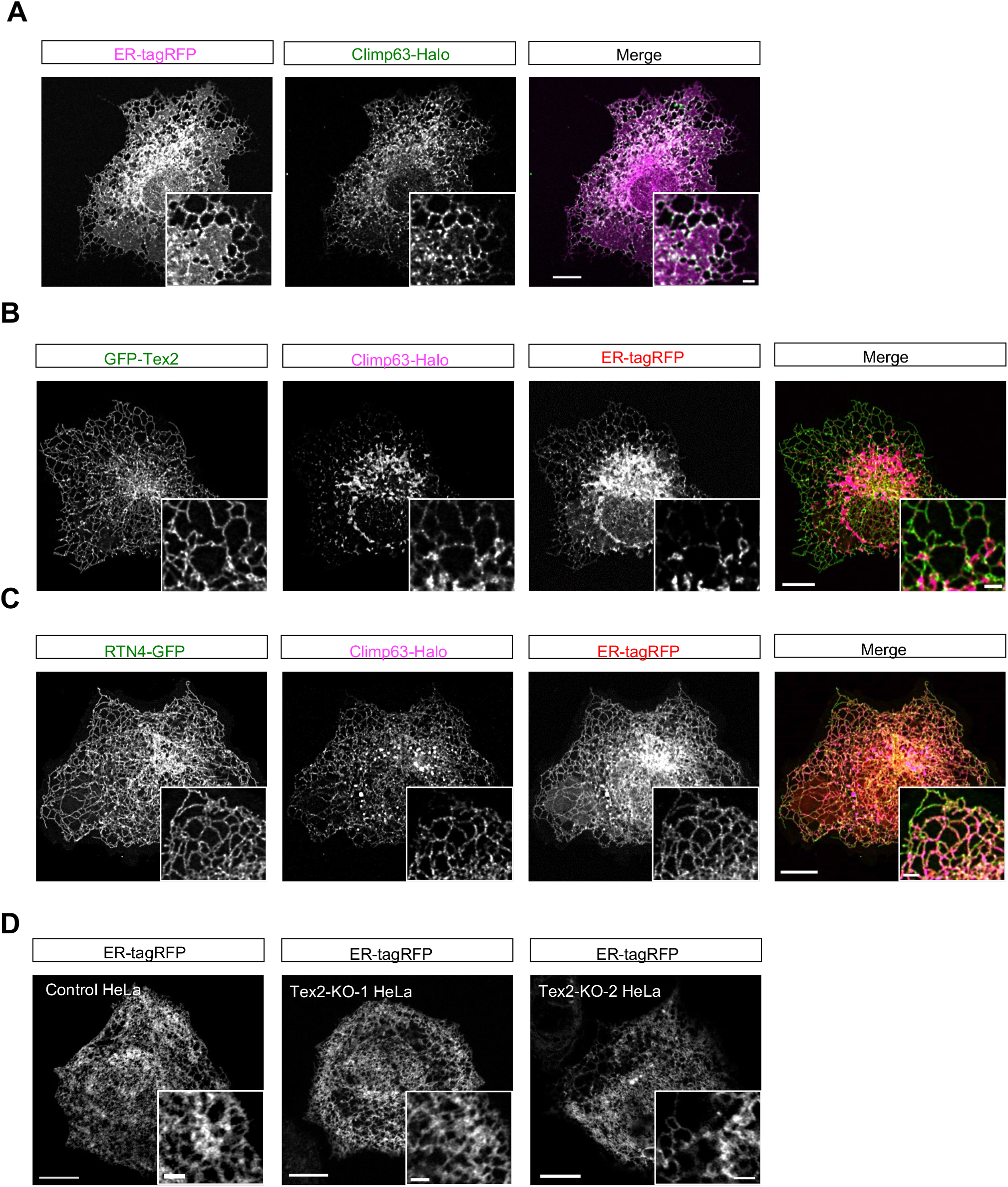
A potential role of Tex2 in tubular ER shaping. **A-C**. Representative images of COS7 cells expressing Climp63-Halo alone (magenta; **A**), Climp63-Halo (magenta) and GFP-Tex2 (green) (**B**), or Climp63-Halo (magenta) and RTN4-GFP (green) (**C**) with insets at cell periphery. **D**. Representative images of control, or two HeLa Tex2-KO clones expressing ER-tagRFP (magenta) with insets at cell periphery. Scale bar, 10μm in the whole cell images and 2μm in the insets in (A-D).

**Fig. S5.**
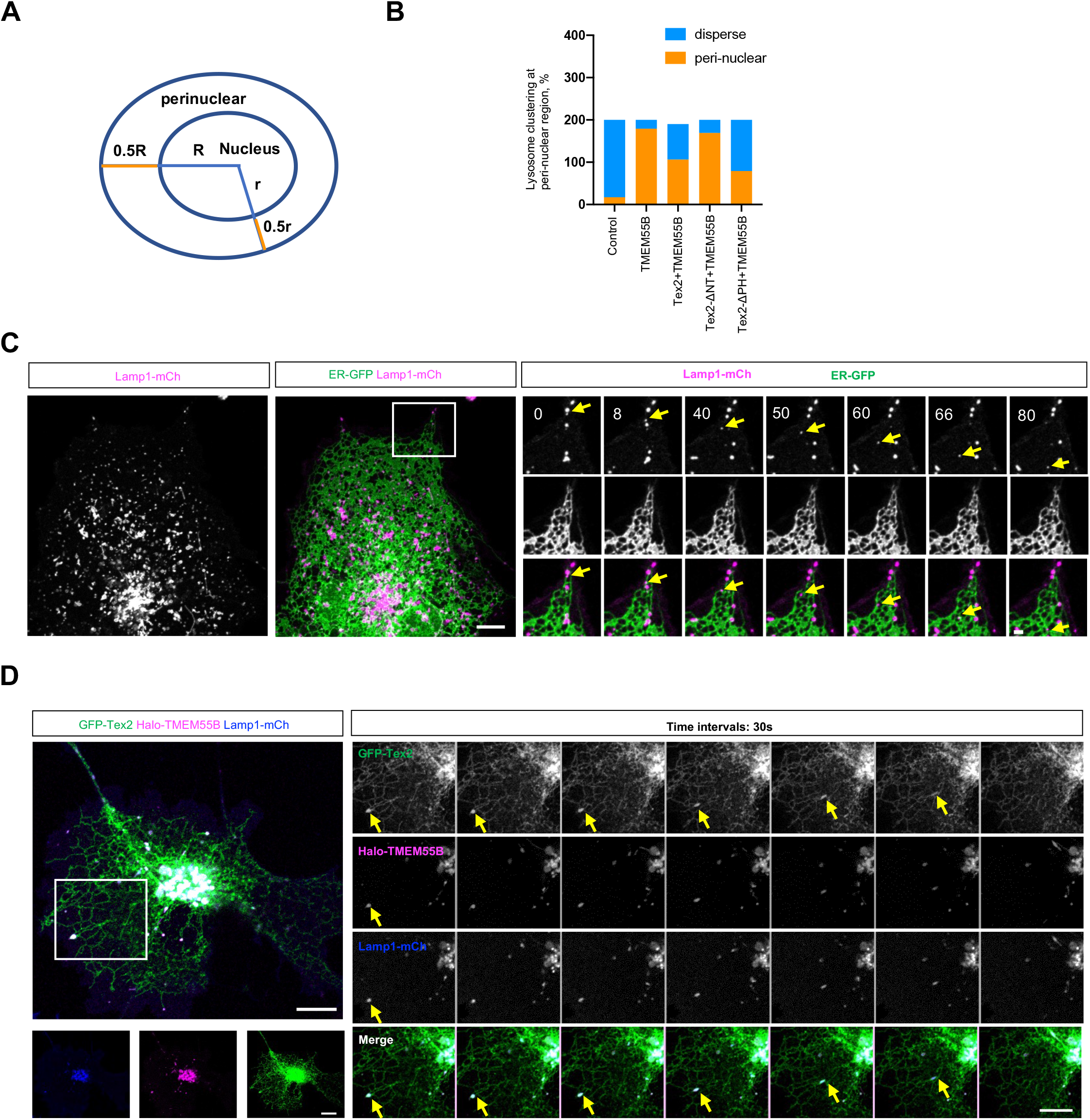
A potential role of Tex2-TMEM55B interaction in retrograde transport of LE/lys. **A.** Schematic cartoon showing the definition of peri-nuclear regions. **B.** Percentage of cells with lysosome clustering at peri-nuclear region in control (200 cells), TMEM55B overexpression (200 cells), co-overexpression of TMEM55B and Tex2 (184 cells), co-overexpression of TMEM55B and Tex2-ΔNT (1-514) (200 cells), and co-overexpression of TMEM55B and Tex2-ΔPH (200 cells) in more than 3 independent assays. Ordinary one-way ANOVA with Tukey’s multiple comparisons test. Mean ± SD. **C.** Representative images of a live COS7 cell expressing ER-GFP (green), and Lamp1-mCh (magenta) with time-lapse images of a boxed region showing on the right. Yellow arrows denote one LE/lys undergoing retrograde transport with frequent contacts with the ER. **D.** Representative images of a live COS7 cell expressing GFP-Tex2 (green), Halo-TMEM55B (magenta), and Lamp1-mCh (blue) with time-lapse images of a boxed region showing on the right. Yellow arrows denote one LE/lys undergoing retrograde transport along with the Tex2-labeled ER. Scale bar, 10μm in the whole cell images and 2μm in the insets in (C, D).

**Fig. S6.**
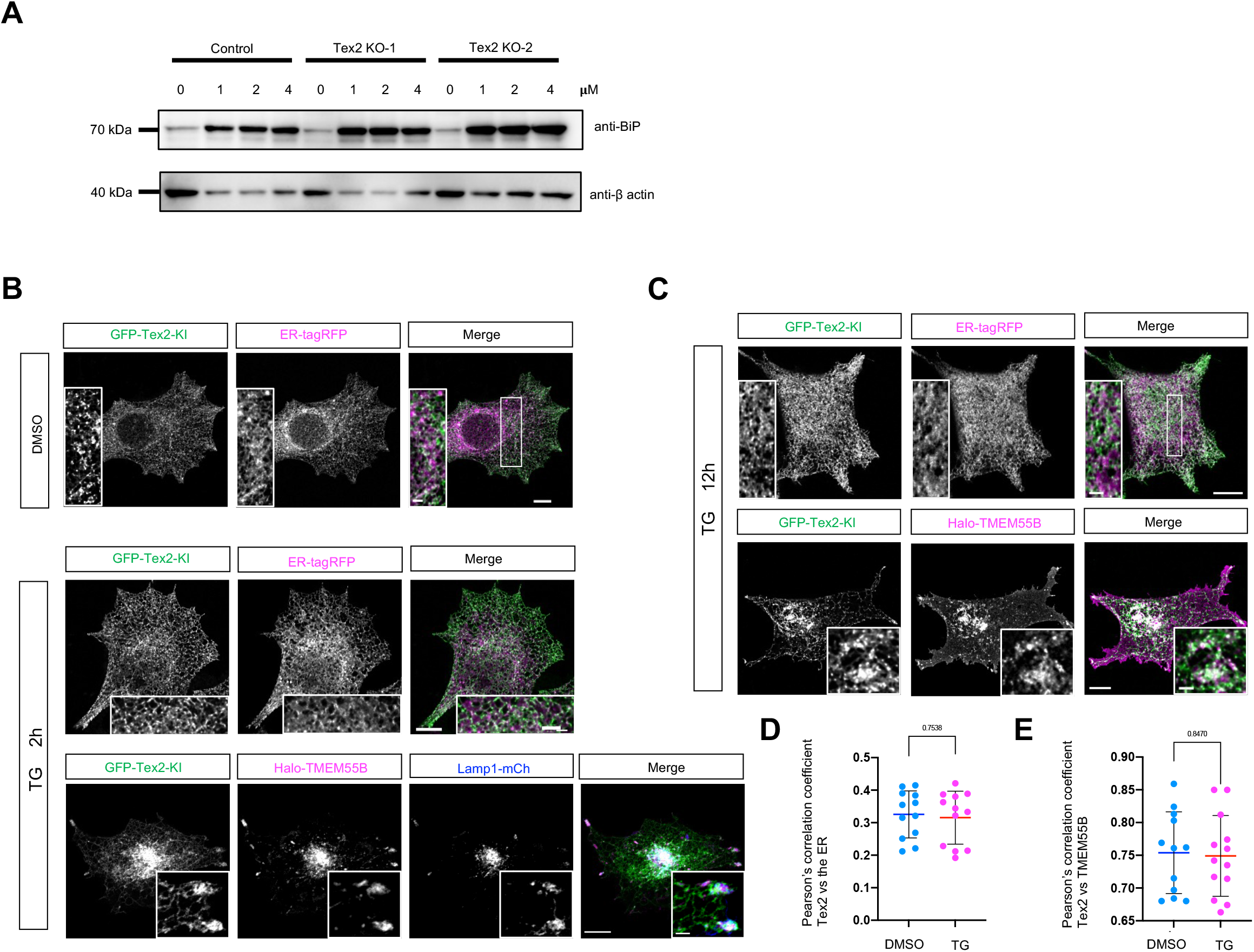
Tex2 may not be directly involved in TG-induced ER stress in mammalian cells. **A.** Western blots demonstrate the increase in BiP level in control or two Tex2-KO clones upon TG stimulation for 12h. **B.** Representative images of a GFP-Tex2-KI (green) cell expressing ER-tagRFP (magenta) upon DMSO (top), TG (2h; middle), or expressing Halo-TMEM55B (magenta) and Lamp1-mCh (blue) upon TG treatments (2h; bottom) with insets. **C.** Representative images of a GFP-Tex2-KI (green) cell expressing ER-tagRFP (magenta; top) or Halo-TMEM55B (magenta; bottom) upon TG treatment for 12h with insets. **D.** Pearson’s correlation coefficient of GFP-Tex2 vs ER-tagRFP in DMSO (12 cells) or TG-treated cell (12 cells). Two-tailed unpaired student t-test. Mean ± SD. **E.** Pearson’s correlation coefficient of GFP-Tex2 vs Halo-TMEM55B in DMSO (12 cells) or TG-treated cell (13 cells). Two-tailed unpaired student t-test. Mean ± SD. Scale bar, 10μm in the whole cell images and 2μm in the insets in (B, C).

**Fig. S7.**
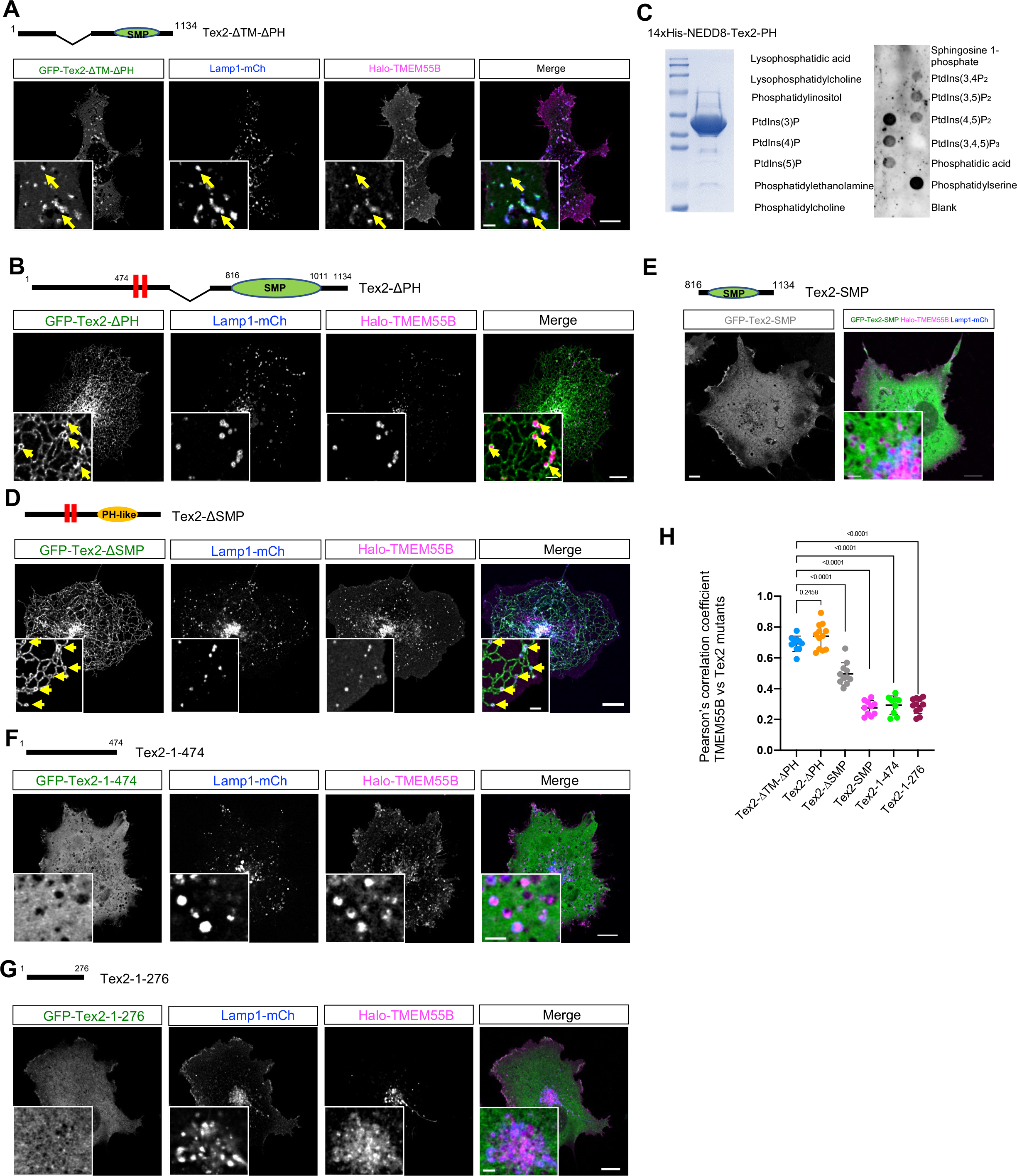
Neither the PH-like domain nor the SMP domain is responsible for interacting with TMEM55B. **A-B**. Representative images of a COS7 cell expressing either Tex2-ΔTM-ΔPH (green; **A**), or Tex2-ΔPH (green; **B**), along with Lamp1-mCh (blue) and Halo-TMEM55B (magenta) with insets. Yellow arrows indicate Tex2 enrichments at TMEM55B-positive LE/lys. **C**. PIPs strip assay of purified Tex2-PH. Left: Coomassie blue staining of purified Tex2-PH; right: PIPs strip blots. **D**. Representative images of a COS7 cell expressing Tex2-ΔSMP (green) and Lamp1-mCh (blue) and Halo-TMEM55B (magenta) with insets. Yellow arrows indicate Tex2 proteins enrichment at TMEM55B-positive LE/lys. **E**. Representative images of a COS7 cell expressing Tex2-SMP alone (gray; left) or Tex2-SMP along with Lamp1-mCh (blue) and Halo-TMEM55B (magenta) with an inset (right). **F-G**. Representative images of a COS7 cell expressing Tex2-1-474 (green; **F**) or Tex2-1-276 (green; **G**), along with Lamp1-mCh (blue) and Halo-TMEM55B (magenta) with insets. **H**. Pearson’s correlation coefficient of either Tex2-ΔTM-ΔPH (10 cells), Tex2-ΔPH (10 cells), Tex2-ΔSMP (11 cells), Tex2-SMP (9 cells), Tex2-1-474 (9 cells), or Tex2-1-276 (11 cells) vs Halo-TMEM55B in 3 independent assays. Ordinary one-way ANOVA with Tukey’s multiple comparisons test. Mean ± SD. **I**. As in Fig. 5H, pulldown assays using GFP-Tex2 bound on GFP-Trap beads and purified GST-TMEM55B demonstrate a direct interaction in a PS or ceramide-independent manner. Scale bar, 10μm in the whole cell images and 2μm in the insets in (A, B, D, E, G, & H).

**Fig. S8.**
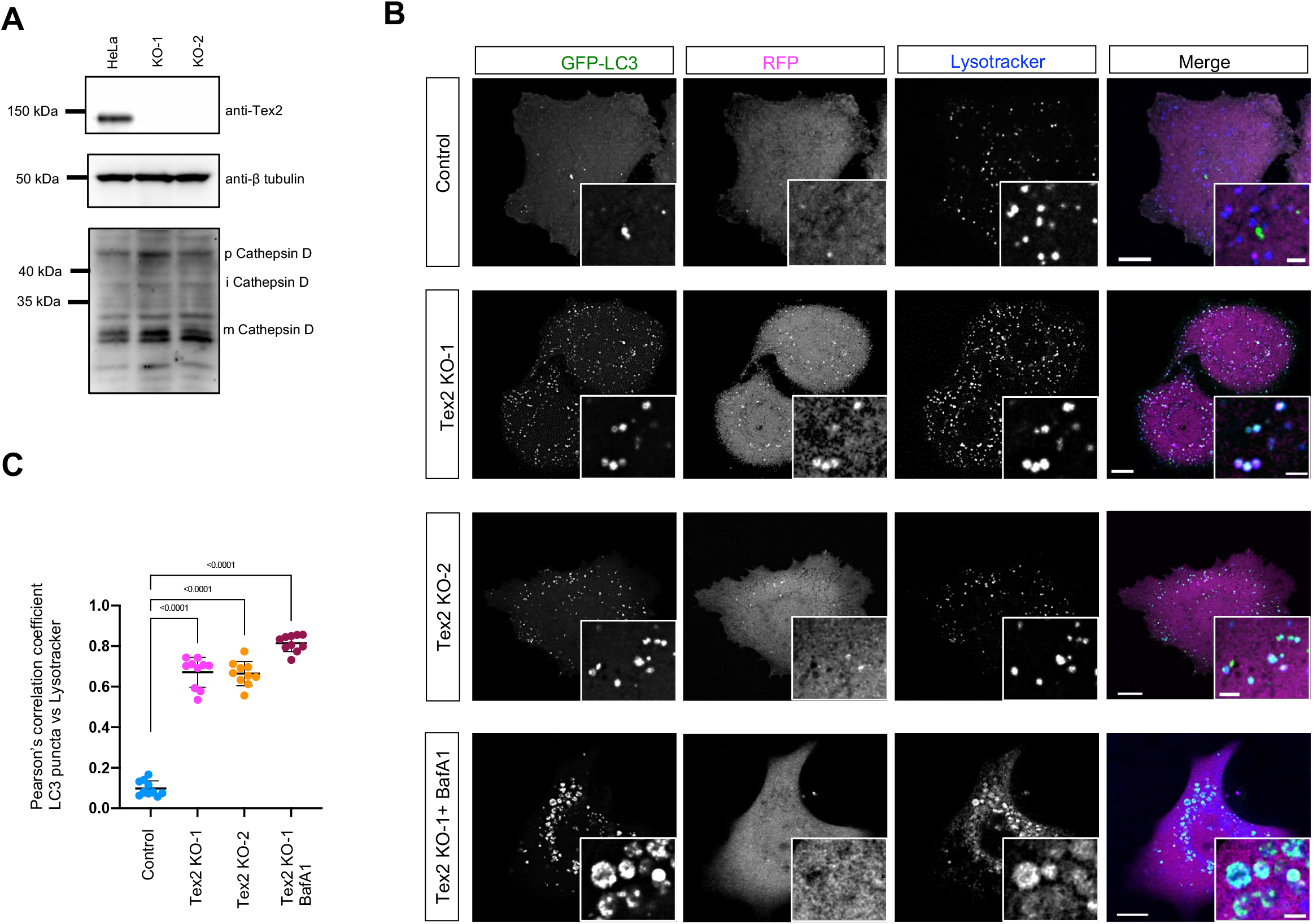
Tex2 KO does not block the autophagosome-lysosome fusion. **A.** Western blots show the level of CathepsinD in control or two Tex2-KO clones. **B.** Representative images of either control (top), two Tex2-KO clones (middle) or Tex2-KO-1 cells treated with BafA1 expressing GFP-LC3-RFP were labeled with Lysotracker (blue) with insets. **C.** Pearson’s correlation coefficient of GFP-LC3 vs Lysotracker in control (10 cells), Tex2-KO-1 (10 cells), Tex2-KO-2 (10 cells), or Tex2-KO-1 treated with BafA1 (10 cells) in 3 independent assays. Ordinary one-way ANOVA with Tukey’s multiple comparisons test. Mean ± SD. Scale bar, 10μm in the whole cell images and 2μm in the insets in (B).

**Fig. S9.**
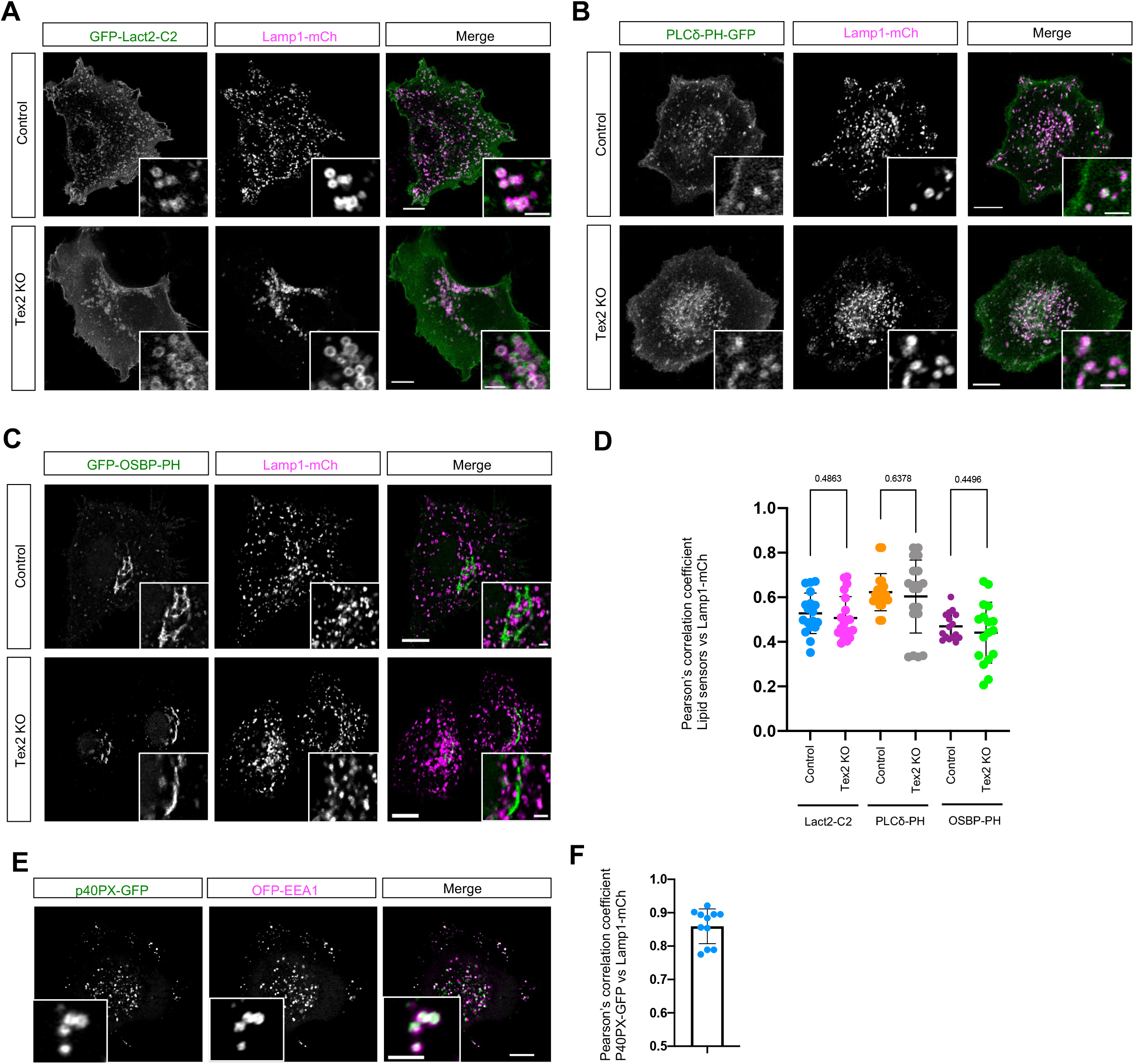
PS, PI4P, and PI(4,5)P2 in Tex2 KO cells. **A-C**. Representative images of either control (top) or Tex2-KO-1 (bottom) cells expressing either GFP-Lact2-C2 (green; PS sensor; **A**), PLCδ-PH-GFP (green; PI45P2 sensor; **B**), or GFP-OSBP-PH (green; PI4P sensor; **C**) with insets. **D.** Pearson’s correlation coefficient of either GFP-Lact2-C2 (18 control; 20 Tex2-KO cells), PLCδ-PH-GFP (20 control; 20 Tex2-KO cells), GFP-OSBP-PH (16 control; 17 Tex2-KO cells), or vs Lamp1-mCh. Two-tailed unpaired student t-test. Mean ± SD. **E.** Representative images of a control cell expressing p40PX-GFP (green; a PI3P sensor) and OFP-EEA1 (magenta) with insets. **F.** Pearson’s correlation coefficient of p40PX-GFP vs OFP-EEA1 (11 cells). Mean ± SD. Scale bar, 10μm in the whole cell images and 2μm in the insets in (A-C).

## Notes

### Competing Interest Statement

The authors have declared no competing interest.

## Reference

Anderson, D.J., and M.W. Hetzer. 2008. Reshaping of the endoplasmic reticulum limits the rate for nuclear envelope formation. J Cell Biol. 182:911–924.

Balla, A., G. Tuymetova, M. Barshishat, M. Geiszt, and T. Balla. 2002. Characterization of type II phosphatidylinositol 4-kinase isoforms reveals association of the enzymes with endosomal vesicular compartments. J Biol Chem. 277:20041–20050.

Baumann, O., and B. Walz. 2001. Endoplasmic reticulum of animal cells and its organization into structural and functional domains. Int Rev Cytol. 205:149–214.

Bian, X., R.W. Klemm, T.Y. Liu, M. Zhang, S. Sun, X. Sui, X. Liu, T.A. Rapoport, and J. Hu. 2011. Structures of the atlastin GTPase provide insight into homotypic fusion of endoplasmic reticulum membranes. Proc Natl Acad Sci U S A. 108:3976–3981.

Bian, X., Y. Saheki, and P. De Camilli. 2018. Ca(2+) releases E-Syt1 autoinhibition to couple ER-plasma membrane tethering with lipid transport. EMBO J. 37:219–234.

Borgese, N., M. Francolini, and E. Snapp. 2006. Endoplasmic reticulum architecture: structures in flux. Curr Opin Cell Biol. 18:358–364.

Chen, F., B. Yan, J. Ren, R. Lyu, Y. Wu, Y. Guo, D. Li, H. Zhang, and J. Hu. 2021. FIT2 organizes lipid droplet biogenesis with ER tubule-forming proteins and septins. J Cell Biol. 220.

Dikic, I., and Z. Elazar. 2018. Mechanism and medical implications of mammalian autophagy. Nat Rev Mol Cell Biol. 19:349–364.

Dong, R., T. Zhu, L. Benedetti, S. Gowrishankar, H. Deng, Y. Cai, X. Wang, K. Shen, and P. De Camilli. 2018. The inositol 5-phosphatase INPP5K participates in the fine control of ER organization. J Cell Biol. 217:3577–3592.

English, A.R., and G.K. Voeltz. 2013. Endoplasmic reticulum structure and interconnections with other organelles. Cold Spring Harb Perspect Biol. 5:a013227.

Friedman, J.R., J.R. Dibenedetto, M. West, A.A. Rowland, and G.K. Voeltz. 2013. Endoplasmic reticulum-endosome contact increases as endosomes traffic and mature. Mol Biol Cell. 24:1030–1040.

Gao, Y., J. Xiong, Q.Z. Chu, and W.K. Ji. 2022. PDZD8-mediated lipid transfer at contacts between the ER and late endosomes/lysosomes is required for neurite outgrowth. J Cell Sci. 135.

Gillooly, D.J., C. Raiborg, and H. Stenmark. 2003. Phosphatidylinositol 3-phosphate is found in microdomains of early endosomes. Histochem Cell Biol. 120:445–453.

Giordano, F., Y. Saheki, O. Idevall-Hagren, S.F. Colombo, M. Pirruccello, I. Milosevic, E.O. Gracheva, S.N. Bagriantsev, N. Borgese, and P. De Camilli. 2013. PI(4,5)P(2)-dependent and Ca(2+)-regulated ER-PM interactions mediated by the extended synaptotagmins. Cell. 153:1494–1509.

Holthuis, J.C., and T.P. Levine. 2005. Lipid traffic: floppy drives and a superhighway. Nat Rev Mol Cell Biol. 6:209–220.

Hoyer, M.J., P.J. Chitwood, C.C. Ebmeier, J.F. Striepen, R.Z. Qi, W.M. Old, and G.K. Voeltz. 2018. A Novel Class of ER Membrane Proteins Regulates ER-Associated Endosome Fission. Cell. 175:254–265 e214.

Jeyasimman, D., B. Ercan, D. Dharmawan, T. Naito, J. Sun, and Y. Saheki. 2021. PDZD-8 and TEX-2 regulate endosomal PI(4,5)P2 homeostasis via lipid transport to promote embryogenesis in C. elegans. Nat Commun. 12:6065.

Ji, W.K., R. Chakrabarti, X. Fan, L. Schoenfeld, S. Strack, and H.N. Higgs. 2017. Receptor-mediated Drp1 oligomerization on endoplasmic reticulum. J Cell Biol. 216:4123–4139.

Jongsma, M.L., I. Berlin, R.H. Wijdeven, L. Janssen, G.M. Janssen, M.A. Garstka, H. Janssen, M. Mensink, P.A. van Veelen, R.M. Spaapen, and J. Neefjes. 2016. An ER-Associated Pathway Defines Endosomal Architecture for Controlled Cargo Transport. Cell. 166:152–166.

Joshi, A.S., H. Zhang, and W.A. Prinz. 2017. Organelle biogenesis in the endoplasmic reticulum. Nat Cell Biol. 19:876–882.

Kaizuka, T., H. Morishita, Y. Hama, S. Tsukamoto, T. Matsui, Y. Toyota, A. Kodama, T. Ishihara, T. Mizushima, and N. Mizushima. 2016. An Autophagic Flux Probe that Releases an Internal Control. Mol Cell. 64:835–849.

Kanai, F., H. Liu, S.J. Field, H. Akbary, T. Matsuo, G.E. Brown, L.C. Cantley, and M.B. Yaffe. 2001. The PX domains of p47phox and p40phox bind to lipid products of PI(3)K. Nat Cell Biol. 3:675–678.

Kornmann, B., E. Currie, S.R. Collins, M. Schuldiner, J. Nunnari, J.S. Weissman, and P. Walter. 2009. An ER-mitochondria tethering complex revealed by a synthetic biology screen. Science. 325:477–481.

Kumar, D., B. Golchoubian, I. Belevich, E. Jokitalo, and A.L. Schlaitz. 2019. REEP3 and REEP4 determine the tubular morphology of the endoplasmic reticulum during mitosis. Mol Biol Cell. 30:1377–1389.

Lebiedzinska, M., G. Szabadkai, A.W. Jones, J. Duszynski, and M.R. Wieckowski. 2009. Interactions between the endoplasmic reticulum, mitochondria, plasma membrane and other subcellular organelles. Int J Biochem Cell Biol. 41:1805–1816.

Lees, J.A., M. Messa, E.W. Sun, H. Wheeler, F. Torta, M.R. Wenk, P. De Camilli, and K.M. Reinisch. 2017. Lipid transport by TMEM24 at ER-plasma membrane contacts regulates pulsatile insulin secretion. Science. 355.

Lemmon, M.A. 2007. Pleckstrin homology (PH) domains and phosphoinositides. Biochem Soc Symp:81–93.

Levine, T. 2004. Short-range intracellular trafficking of small molecules across endoplasmic reticulum junctions. Trends Cell Biol. 14:483–490.

Levine, T. 2005. A new way for sterols to walk on water. Mol Cell. 19:722–723.

Levine, T., and C. Rabouille. 2005. Endoplasmic reticulum: one continuous network compartmentalized by extrinsic cues. Curr Opin Cell Biol. 17:362–368.

Levine, T.P., and S. Munro. 2002. Targeting of Golgi-specific pleckstrin homology domains involves both PtdIns 4-kinase-dependent and -independent components. Curr Biol. 12:695–704.

Li, P., J.A. Lees, C.P. Lusk, and K.M. Reinisch. 2020. Cryo-EM reconstruction of a VPS13 fragment reveals a long groove to channel lipids between membranes. J Cell Biol. 219.

Liu, L.K., V. Choudhary, A. Toulmay, and W.A. Prinz. 2017. An inducible ER-Golgi tether facilitates ceramide transport to alleviate lipotoxicity. J Cell Biol. 216:131–147.

Marat, A.L., and V. Haucke. 2016. Phosphatidylinositol 3-phosphates-at the interface between cell signalling and membrane traffic. EMBO J. 35:561–579.

Moreno, R.D., and C.P. Alvarado. 2006. The mammalian acrosome as a secretory lysosome: new and old evidence. Mol Reprod Dev. 73:1430–1434.

Prinz, W.A., A. Toulmay, and T. Balla. 2020. The functional universe of membrane contact sites. Nat Rev Mol Cell Biol. 21:7–24.

Raiborg, C., E.M. Wenzel, N.M. Pedersen, H. Olsvik, K.O. Schink, S.W. Schultz, M. Vietri, V. Nisi, C. Bucci, A. Brech, T. Johansen, and H. Stenmark. 2015. Repeated ER-endosome contacts promote endosome translocation and neurite outgrowth. Nature. 520:234–238.

Rocha, N., C. Kuijl, R. van der Kant, L. Janssen, D. Houben, H. Janssen, W. Zwart, and J. Neefjes. 2009. Cholesterol sensor ORP1L contacts the ER protein VAP to control Rab7-RILP-p150 Glued and late endosome positioning. J Cell Biol. 185:1209–1225.

Rowland, A.A., P.J. Chitwood, M.J. Phillips, and G.K. Voeltz. 2014. ER contact sites define the position and timing of endosome fission. Cell. 159:1027–1041.

Schauder, C.M., X. Wu, Y. Saheki, P. Narayanaswamy, F. Torta, M.R. Wenk, P. De Camilli, and K.M. Reinisch. 2014. Structure of a lipid-bound extended synaptotagmin indicates a role in lipid transfer. Nature. 510:552–555.

Shibata, Y., T. Shemesh, W.A. Prinz, A.F. Palazzo, M.M. Kozlov, and T.A. Rapoport. 2010. Mechanisms determining the morphology of the peripheral ER. Cell. 143:774–788.

Stauffer, T.P., S. Ahn, and T. Meyer. 1998. Receptor-induced transient reduction in plasma membrane PtdIns(4,5)P2 concentration monitored in living cells. Curr Biol. 8:343–346.

Ugur, B., W. Hancock-Cerutti, M. Leonzino, and P. De Camilli. 2020. Role of VPS13, a protein with similarity to ATG2, in physiology and disease. Curr Opin Genet Dev. 65:61–68.

Ungewickell, A., C. Hugge, M. Kisseleva, S.C. Chang, J. Zou, Y. Feng, E.E. Galyov, M. Wilson, and P.W. Majerus. 2005. The identification and characterization of two phosphatidylinositol-4,5-bisphosphate 4-phosphatases. Proc Natl Acad Sci U S A. 102:18854–18859.

Voeltz, G.K., W.A. Prinz, Y. Shibata, J.M. Rist, and T.A. Rapoport. 2006. A class of membrane proteins shaping the tubular endoplasmic reticulum. Cell. 124:573–586.

Wang, S., H. Tukachinsky, F.B. Romano, and T.A. Rapoport. 2016. Cooperation of the ER-shaping proteins atlastin, lunapark, and reticulons to generate a tubular membrane network. Elife. 5.

Willett, R., J.A. Martina, J.P. Zewe, R. Wills, G.R.V. Hammond, and R. Puertollano. 2017. TFEB regulates lysosomal positioning by modulating TMEM55B expression and JIP4 recruitment to lysosomes. Nat Commun. 8:1580.

Wong, L.H., A. Copic, and T.P. Levine. 2017. Advances on the Transfer of Lipids by Lipid Transfer Proteins. Trends Biochem Sci. 42:516–530.

Wong, L.H., A.T. Gatta, and T.P. Levine. 2019. Lipid transfer proteins: the lipid commute via shuttles, bridges and tubes. Nat Rev Mol Cell Biol. 20:85–101.

Wu, H., and G.K. Voeltz. 2021. Reticulon-3 Promotes Endosome Maturation at ER Membrane Contact Sites. Dev Cell. 56:52–66 e57.

Yamamoto, Y., A. Yoshida, N. Miyazaki, K. Iwasaki, and T. Sakisaka. 2014. Arl6IP1 has the ability to shape the mammalian ER membrane in a reticulon-like fashion. Biochem J. 458:69–79.

Yeung, T., G.E. Gilbert, J. Shi, J. Silvius, A. Kapus, and S. Grinstein. 2008. Membrane phosphatidylserine regulates surface charge and protein localization. Science. 319:210–213.

Zajac, A.L., Y.E. Goldman, E.L. Holzbaur, and E.M. Ostap. 2013. Local cytoskeletal and organelle interactions impact molecular-motor-driven early endosomal trafficking. Curr Biol. 23:1173–1180.

Zhou, Q., J. Li, H. Yu, Y. Zhai, Z. Gao, Y. Liu, X. Pang, L. Zhang, K. Schulten, F. Sun, and C. Chen. 2014. Molecular insights into the membrane-associated phosphatidylinositol 4-kinase IIalpha. Nat Commun. 5:3552.

